# Lipogenic Lung Fibroblast-derived Extracellular Vesicles Mitigate Cigarette Smoke-Induced Chronic Obstructive Pulmonary Disease Pathologies through LAT1-mediated Alveolar Type II Cell Restoration

**DOI:** 10.1101/2024.06.17.587086

**Authors:** Shota Fujimoto, Yuta Hirano, Naoaki Watanabe, Sachi Matsubayashi, Shun Inukai, Saiko Nishioka, Masahiro Yoshida, Saburo Ito, Shunsuke Minagawa, Hiromichi Hara, Takashi Ohtsuka, Pattama Wiriyasermkul, Shushi Nagamori, Kazuyoshi Kuwano, Jun Araya, Yu Fujita

**Affiliations:** Division of Respiratory Diseases, Department of Internal Medicine, The Jikei University School of Medicine, Tokyo, Japan; Division of Thoracic Surgery, Department of Surgery, The Jikei University School of Medicine, Tokyo, Japan; Center for SI Medical Research, The Jikei University School of Medicine, Tokyo, Japan; Department of Biological Chemistry and Food Science, Faculty of Agriculture, Iwate University, Iwate, Japan; Department of Laboratory Medicine, The Jikei University School of Medicine, Tokyo, Japan; Division of Next-Generation Drug Development, Research Center for Medical Sciences, The Jikei University School of Medicine, Tokyo, Japan; Center for Exosome Medical Research, The Jikei University School of Medicine, Tokyo, Japan

**Author notes:** Address correspondence to: Yu Fujita, The Jikei University School of Medicine, Nishi-shimbashi 3-25-8, Minato-ku, Tokyo, 105-8461, Japan.

## Abstract

Emerging research has revealed specific cellular aberrations in Chronic Obstructive Pulmonary Disease (COPD), with a particular focus on alveolar type 2 (AT2) cells, which play a pivotal role in the restoration of damaged lung tissue and promotion of normal cellular differentiation. Lipofibroblasts (LipoFBs), which are stromal fibroblasts that house lipid droplets, have been identified in close proximity to AT2 cells and have been demonstrated to support AT2 function. In this study, we present a comprehensive investigation into the therapeutic potential of extracellular vesicles (EVs) derived from LipoFBs (LipoFB-EVs) in COPD treatment. They effectively mitigate key COPD pathologies such as cellular senescence and inflammatory responses in lung epithelial cells. This is achieved by reducing reactive oxygen species (ROS) levels and modulating DNA damage response pathways. Moreover, LipoFB-EVs demonstrate antifibrotic properties by inhibiting TGF-β-induced myofibroblast differentiation, surpassing conventional antifibrotic drugs. They also aid in restoring impaired AT2 stem cells, which are crucial for lung homeostasis, by enhancing their viability, colony-forming ability, and proliferation. Furthermore, we identify the presence of L-type amino acid transporter 1 (LAT1) within LipoFB-EVs, which mediates amino acid uptake, particularly leucine transport, and contributes to the restoration of AT2 cell dysfunction. Importantly, the administration of LipoFB-EVs in murine models of COPD resulted in significant improvements in airway inflammation, remodeling, obstruction, cellular senescence, and alveolar emphysema induced by both short- and long-term CS exposure. Overall, our findings highlight the therapeutic potential of LipoFB-EVs as a novel regenerative therapy for COPD, offering promising avenues for future clinical interventions.

## Introduction

Chronic Obstructive Pulmonary Disease (COPD) is a prevalent respiratory disease characterized by progressive and irreversible airflow limitation resulting from increased tissue degradation and impaired tissue repair mechanisms^1^. Existing therapeutic approaches have shown limited efficacy in halting disease progression, emphasizing the pressing need for innovative pharmacological interventions aimed at reactivating lung repair processes^2^. Recent advances in organoid culture techniques and single-cell transcriptome analyses have shed light on the intricate molecular landscape of COPD pathogenesis^3–6^. These studies revealed inflammatory perturbations in alveolar type 2 (AT2) cells, disturbances in protein homeostasis, cellular senescence, and functional aberrations during the healing process, which collectively contribute to the loss of AT2 stemness. This finding underscores the potential for enhancing AT2 stemness as a promising avenue for promoting the normal differentiation of alveolar type 1 (AT1) cells and rejuvenating damaged lung parenchyma, offering prospects for regenerative therapy in COPD. Nonetheless, the present arsenal of drugs targeting peripheral airway fibrosis and alveolar structural damage, which are the primary pathogenic drivers of COPD, have demonstrated limited antifibrotic and lung rejuvenation effects. Consequently, there exists an urgent imperative for the development of novel therapies dedicated to lung regeneration, addressing this unmet medical need.

Lung fibroblasts (LFs), an integral component of lung tissue, exhibit a fascinating spectrum of functional diversity. Within this population, two prominent subtypes contribute to this diversity: lipogenic and myogenic subtypes^7^. Lipofibroblasts (LipoFBs), characterized by the presence of lipid droplets, reside within the lung stroma in close proximity to AT2 cells^8^. Functionally, LipoFBs, in contrast to myofibroblasts (MyoFBs), display intriguing antifibrotic properties. Moreover, these cells actively participate in the differentiation and sustenance of AT2 cell function, acting as niche cells for epithelial stem cells^9^. In the alveolar microenvironment, LipoFBs play a role in lung repair and regeneration following injury and fibrosis^10,11^. Furthermore, dysregulation of LipoFB function in the lungs can have implications for alveolar disorders, including idiopathic pulmonary fibrosis (IPF)^7,11^. However, the precise functions and contributions of LipoFBs in COPD have not yet been fully elucidated.

In IPF, the lung tissue demonstrates a shift from LipoFBs to MyoFBs, implying that the conversion of LipoFBs into MyoFBs plays a pivotal role in the development of lung fibrosis^9^. Previous reports indicate that LipoFBs can be induced to differentiate by metabolic reprogramming of the glycolytic and fatty acid systems in a cell culture system using the diabetes drug, Metformin, or the peroxisome proliferator-activated receptor gamma (PPARγ) activator, Rosiglitazone, and that the induction of MyoFBs to LipoFBs may be an antifibrotic therapy^9,12^. The mechanism is thought to involve the enhancement of extracellular matrix degradation and the suppression of TGF-β by regulating the intracellular glycolytic system and fatty acid metabolism. However, studies on LipoFB-induced differentiation that have been demonstrated to date have used high concentrations of drugs, which has been a huge barrier to the clinical application of the antifibrotic effects of metabolic reprogramming and the effects of LipoFB-secreted factors on AT2 cells to maintain function. Therefore, we focused on metabolically reprogrammed LF-derived extracellular vesicles (EVs), which are an important tool for cell-to-cell communication, in the development of new lung regeneration therapies.

EVs can be categorized into several types based on their biogenesis and size, including exosomes (30-150 nm), microvesicles (100-1000 nm), and apoptotic bodies (800-5000 nm)^13^. EVs are enveloped in a lipid bilayer membrane and are ubiquitously secreted by all cells. These vesicles exhibited remarkable stability in various bodily fluids. They contain nucleic acids such as proteins and microRNAs (miRNAs), which reflect the internal environment of secretory cells and transfer them between cells. Understanding cell-to-cell communication via EVs has led to the development of disease pathogenesis, biomarkers, and, in the past few years, therapeutic applications^14^. Recent evidence that the efficacy of cellular therapies, such as mesenchymal stem cell (MSC)-based therapies for the treatment of refractory diseases, may be mediated by humoral factors, and in part by EVs, has challenged researchers worldwide to apply MSC-derived EVs in clinical practice, including chronic lung diseases^15^. As therapeutic agents, EVs are characterized by their multi-targeted efficacy owing to the inclusion of diverse components, and they have advantages over conventional respiratory disease drug discovery. While MSCs and various other candidates, such as bronchial epithelium^16^ and lung spheroid cells^17^, show promise in the context of EV-based therapy for patients with IPF ^18^, the landscape of COPD lacks compelling EV-based therapeutic breakthroughs. Within the intricate milieu of chronic airway inflammation, fibrosis, and impaired lung repair, COPD presents a challenging arena with a notable scarcity of innovative EV-based treatment candidates. Considering that damage to AT2 cells is a critical step in COPD pathogenesis, it is plausible that EVs released by LipoFBs possess antifibrotic potential and the capacity to impede the progression of COPD pathology by influencing AT2 stemness, potentially serving as niche-like entities. In this study, we aimed to elucidate whether LipoFB-derived EVs could address small airway remodeling and alveolar structural destruction in COPD, offering therapeutic benefits, such as antifibrotic and anti-aging effects and AT2 stem cell restoration.

## Methods and Materials Single-cell RNA sequencing

Raw sequencing data were converted into a cell expression matrix using the software, Cell Ranger (version 6.1.2, 10x Genomics), followed by quality filtering, data normalization, assessment, and correction of batch effects using Seurat (version 5.0.1)^19^. Briefly, low-quality cells were filtered based on the parameters ’nFeature_RNA > 500 & nFeature_RNA < 8000 & percent.mt < 20.’ A meta-single-cell cohort, including 116 patients and 601,887 cells, was utilized for subsequent analyses (**sTable 1**). Subsequently, the metadata was normalized, scaled, and subjected to principal component analysis (PCA) using Seurat functions. All single-cell datasets were integrated, and batch effects were adjusted using Harmony (version 1.2.0)^20^. Cell clustering was performed using the ’FindNeighbors’ and ’FindClusters’ functions from Seurat, and the dimensionality-reduced cell clustering was visualized using Uniform Manifold Approximation and Projection (UMAP) plots generated by the ’runUMAP’ function. Cell type annotation was based on published papers and SingleR (version 2.4.1)^21,22^. Annotated cell clusters were validated using known cell marker lists. Differential expression analysis was conducted using the software, Model-based Analysis of Single-cell Transcriptomics (MAST) (version 1.28.0)^23^. The metabolic activity of alveolar epithelial type 2 cells (AT2 cells) was quantified at the single-cell level using scMetabolism, version 0.2.1, an R package, as demonstrated previously ^24^.

### Ingenuity Pathway Analysis

Genes with a fold change greater than 1.5 and an adjusted p-value less than 0.05 were considered differentially expressed and investigated using the Ingenuity Pathway Analysis (IPA) core pathway (Qiagen Ingenuity Systems).

## Cell culture and clinical samples

All human lung tissue samples were obtained from pneumonectomy and lobectomy specimens at The Jikei University School of Medicine. LFs and human bronchial epithelial cells (HBECs) were isolated from explant cultures in the same way as previously described ^25^. Briefly, fibroblasts outgrown from lung fragments were cultured in Dulbecco’s modified Eagle’s medium (DMEM) (Gibco, 11965092) with a 10% FBS (Gibco, 26140079) and penicillin-streptomycin (Gibco, 15140122) at 37 [in 5% CO2. LFs were serially passaged when the cells reached ∼80–100% confluence and actively proliferating. These cells were used for experiments until passage five. HBECs were isolated from normal airway tissue (1^st^ through 4^th^ order bronchi) by protease treatment. Freshly isolated HBECs were plated onto rat tail collagen type I-coated (10 μg/ml) dishes in Bronchial Epithelial Growth Medium (BEGM) (Lonza, CC-3170) containing penicillin-streptomycin and grown at 37°C in 5% CO2. HBECs were used until passage 3. Human AT2 cells were isolated and purified as described previously^26^. Briefly, a piece of lung tissue was mechanically minced and dissociated using enzymes containing DNase I (330 U/mL) (Sigma-Aldrich Cat, 10104159001), Collagenase type I (4500 U/mL) (Gibco, 17100-017), and Dispase I (50 U/mL) (Corning, 354235). Samples were incubated at 37°C for 1 hour with constant horizontal rotation. Cells were washed and passed over 100 μm (Corning, 352360) and 40 μm cell strainer (Corning, 352340), and red blood cells were lysed with red-blood-cell lysis buffer (Miltenyi Biotec, 42574000). The cells were then incubated with HTII-280 antibody (Terrace Biotech, TRB-TB-27AHT2-28) and coupled with anti-mouse IgM-coated magnetic beads (Miltenyi Biotec, 130-047-302) to allow for magnetic bead-based AT2 cell separation. The Miltenyi MACS bead sorting kit was further used to separate AT2 cells according to the manufacturer’s instructions (Miltenyi Biotec, Germany). Subsequently, cells were washed with advanced DMEM/F12 (Gibco, 12634028), counted, and seeded in Matrigel (Corning, 354230). Upon solidification of the Matrigel, serum- and feeder-free medium was added to the cultures. Organoids were incubated in an incubator at 37°C and 5% CO2. Informed consent was obtained from all surgical participants as part of an ongoing research protocol approved by the Ethical Committee of the Jikei University School of Medicine (#20-153 (5443).

## Induction of LipoFB

Primary LFs were maintained in DMEM supplemented with 10% FBS at 37°C and 5% CO2. The next day, the cells were starved (serum-free) for 24 hours and treated with 5 mM Metformin (Tokyo Chemical Industry, M2009) or 100 μM Rosiglitazone (Tokyo Chemical Industry, R0106) for 72 hours.

## Antibodies and reagents

Antibodies used were rabbit anti-p21 (Cell Signaling Technology, 2947), mouse anti-p16 (BD Biosciences, 51-1325GR), mouse anti–α smooth muscle actin (Sigma-Aldrich, A2547), goat anti-type I collagen (SouthernBiotech, 1310-01), mouse anti–fibronectin (Abcam, ab6328), rabbit anti-SFTPC (Proteintech, 10774-1-AP), mouse anti-Ki67 (Invitrogen, 14-5698-82), rabbit anti-phosphohistone H2AFX/H2A.X (Ser139) (Cell Signaling Technology, 2577), mouse anti-β-actin (Santa Cruz Biotechnology, sc-47778), rabbit anti–LAT1 (Cell Signaling Technology, 5347), rabbit anti- 4F2hc/CD98 (Cell Signaling Technology, 13180), mouse anti-CD9 (Santa Cruz Biotechnology, sc-59140), mouse anti-CD63 (BD Pharmingen, 556019), rabbit anti-Caveolin-1 antibody (Cell Signaling Technology, 3238), and mouse anti–Tom20 (Santa Cruz Biotechnology, sc-17764). DAPI (Invitrogen, R37606H342), TGF-beta 1 (R&D Systems, 240-B-010), BCH (Cayman, 15249), Pioglitazone (Cayman, 71745), Ciglitazone (Cayman, 71730), DBcAMP (Cayman, 14408), PTHrP (Prospec, HOR-004), Pemafibrate (Kowa, 770515506) Pravastatin (Cayman, 10010342), Pirfenidone (PFD) (Wako, 518-38561), Nintedanib (NTD) (Tocris, 7049), Dithiothreitol (DTT) (Wako, 040-29223), Cell Counting Kit-8 (CCK-8) (DOJINDO, 343-07623), and collagen type I solution from rat tail (Sigma-Aldrich, C3867) were utilized.

## EV purification and analysis

Following growth to 50-80% confluence in 15 cm-cell culture dishes, LFs for EV preparation were washed with PBS. The culture medium was then replaced with fresh advanced DMEM containing penicillin-streptomycin and 2 mM L-glutamine (Gibco, 35050061). After incubation for 48 h, conditioned medium (CM) was collected and centrifuged at 2,000 g for 10 minutes at 4°C. The supernatant was then passed through a 0.22 μm filter (Millipore, SLGV033NB) to thoroughly remove cellular debris. For EV preparation, the CM was ultracentrifuged at 4°C for 45 minutes at 44,200 rpm using an MLS50 rotor. The resulting pellets were washed in PBS, centrifugated by ultracentrifugation, and resuspended in PBS. Protein concentrations of the putative EV fractions were measured using a Qubit 3.0 Fluorometer (Invitrogen). To determine the size distribution of the EVs, nanoparticle tracking analysis was performed using a Nanosight system (NanoSight NS300) with samples diluted 100-fold with PBS. This system focuses a laser beam through a suspension of the particles of interest. The particles were visualized by light scattering using a conventional optical microscope which was perpendicularly aligned to the beam axis, allowing the collection of light scattered from every particle in the field of view. A 60-second video recording of all events was then used for further analysis using the nanoparticle tracking software. The Brownian motion of each particle was used to calculate its size using the Stokes-Einstein equation.

## Electron microscopy

The isolated EVs were visualized using a phase-contrast transmission electron microscope (Terabase Inc., Okazaki, Japan), which can generate high-contrast images of the nanostructures of soft materials, including biological samples, such as liposomes, viruses, bacteria, and cells, without staining processes that might cause damage to the samples. The natural structure of the sample distributed in solution was observed by preparing the sample using a rapid vitreous ice-embedding method and cryo-phase-contrast transmission electron microscopy.

## RNA extraction and qRT-PCR

Total RNA was extracted from cultured cells using QIAzol and the miRNeasy Mini Kit (Qiagen, 217004), following the manufacturer’s protocol. The purity and concentration of all RNA samples were quantified using NanoDrop One (Thermo Fisher Scientific). Reverse transcription was performed using the PrimeScript RT Reagent Kit (Takara, RR037A). The synthesized cDNA was quantified using the Power SYBR Green PCR Master Mix (Thermo Fisher Scientific, 4368706). qRT-PCR analysis was conducted using primers designed using PrimerBank (https://pga.mgh.harvard.edu/primerbank/). β-actin was utilized for normalization. Relative gene expression was measured using the 2(-Delta C(T)) method. Reactions were performed using a QuantStudioTM 3 Real-Time PCR System (Thermo Fisher Scientific) and details of the primers used in the PCR assay are given in **sTable5**.

## Western blot analysis

HBECs and LFs were lysed in Mammalian Protein Extract Reagent (M-PER) (Thermo Fisher Scientific, 78501) containing a protease inhibitor (Roche, 5892970001), phosphatase inhibitor (Roche, 4906845001), and sample buffer solution (Wako, 198-13282 or 196-16142). AT2 organoids were lysed with the above reagents after Matrigel was dissolved in a cell recovery solution (Corning 354253). The protein concentrations were measured by BCA protein assay (Thermo Fisher Scientific, 23227). Western blots of LAT1 and 4F2hc were prepared by mixing samples with sample buffer containing 100 mM dithiothreitol (DTT) under reducing conditions, or without DTT under non-reducing conditions. For each experiment, equal amounts of total protein were resolved by 4–20% SDS-PAGE. After SDS-PAGE, proteins were transferred to a polyvinylidene difluoride membrane (Millipore, ISEQ00010), and incubation with a specific primary Ab was performed overnight at 4°C. After washing several times with PBS containing Tween 20, the membrane was incubated with anti-rabbit IgG, HRP-linked secondary antibody (Cell Signaling Technology, 7074) or anti-mouse IgG, HRP-linked secondary antibody (Cell Signaling Technology, 7076), followed by chemiluminescence detection (Thermo Fisher Scientific, 34080 and Bio-Rad Laboratories, 170-5061) using the ChemiDoc Touch Imaging System (Bio-Rad Laboratories).

## Immunohistochemical staining

Lung tissue samples for immunohistochemistry were collected from C57BL/6J mice. The tissue samples were fixed in formalin and embedded in paraffin. Following dewaxing and rehydration, heat-induced epitope retrieval was performed by boiling the specimens in 1/200 diluted ImmunoSaver (Nissin EM, Tokyo, Japan) at 98°C for 45 min. The endogenous peroxidase activity was inactivated by treating the specimens with 3% H2O2 at room temperature for 10 min. The specimens were treated with 0.1% Triton X-100 for tissue permeabilization. After treatment with a Protein Block Serum-Free blocking reagent (DAKO, Code X0909) at room temperature for 30 min, the specimens were incubated with primary antibodies, like anti-mouse p16 (Abcam, ab54210) and anti-rabbit LAT1 (Trans genic, KE026) at 4°C overnight. The slides were then washed thrice with PBS/ 0.1% Tween-20 and incubated with Dako-labeled polymer-horseradish peroxidase for 1 hour at room temperature. Following this, the sections were incubated with DAB chromogen and counterstained with hematoxylin. The staining intensity of p16 and LAT1 was scored on a four-point scale from 0 (no staining) to 3 (the strongest staining). The intensity scores were multiplied by the percentage of positively stained epithelial cells to generate the IHC score (maximum score, 300).

## Immunofluorescence staining

For immunofluorescence staining of cultured cells, cells were washed thrice with PBS, fixed with 4% paraformaldehyde (Wako, 163-20145), and incubated with primary antibodies containing 0.1% BSA overnight at 4 C. They were then incubated with Alexa Fluor fluorescent secondary antibodies containing 0.1% BSA for 1 hour at room temperature. AT2 organoids were fixed and stained after Matrigel was dissolved in a cell recovery solution. Immunofluorescence staining of paraffin-embedded lung tissues was performed as previously described with minor modifications^16^. Neutral lipids were stained with LipidTOX (1:200) (Invitrogen, H34475). DAPI staining was performed immediately before imaging. Staining was observed using fluorescence microscopy (BZ-X810, Keyence) or confocal microscopy (LSM 880, ZEISS).

## Preparation of cigarette smoke extract

For the preparation of cigarette smoke extract (CSE), approximately 30–50 ml of cigarette smoke was drawn into a syringe and bubbled into sterile PBS in 15-ml BD falcon tubes. One cigarette was used to prepare 10 ml of the solution. To remove insoluble particles, the CSE solution was passed through a 0.22 μm filter (Millipore, SLGS033SS) and was designated as a 100% CSE solution.

## SA-β-Gal Staining

Senescence-associated β-galactosidase (SA-β-Gal) staining was performed using HBECs grown on 24-well culture plates according to the manufacturer’s instructions (Sigma, CS0030).

## Flow cytometry

The cultured fibroblasts were washed with and resuspended in PBS containing LipidTOX (1:200). Following incubation for 30 min, the cells were subjected to flow cytometry using MACS Quant (Miltenyi Biotec). The data was analyzed using the software, FlowJo (FlowJo, LLC).

## Measurement of reactive oxygen species production

HBECs, at a density of 5 × 10^3^ per well, were seeded in a 96-well microplate (Thermo Fisher Scientific, 237105). CM-H2DCFDA was used to measure the total cellular reactive oxygen species (ROS) levels according to the manufacturer’s instructions (Invitrogen, C6827). After incubation with DCFH-DA (100 μM) for 30 minutes at 37°C, the fluorescence of DCF was measured at an excitation wavelength of 485 nm and an emission wavelength of 535 nm by a fluorescence microplate reader (Thermo Scientific, Varioskan LUX).

## Animal studies

Animal experiments were performed in compliance with the guidelines of the Institute for Laboratory Animal Research at the Jikei University School of Medicine (Number: 2018-071). C57BL/6J mice (CLEA Japan INC, Tokyo, Japan) were used for all the experiments. For the chronic cigarette smoke exposure mouse model, 8-week-old mice were exposed to cigarette smoke using a whole-body exposure system (SCIREQ“InExpose”) in a barrier facility. Exposure to cigarette smoke at a total suspended particle (TSP) concentration of 200 mg/m^3^ was carried out using University of Kentucky 3R4F research cigarettes for six days a week over six months. During the final 12 weeks of exposure, sex-matched mice received intratracheal administration of PBS, LF-EVs, or LipoFB(R)-EVs (1.0×10^9^ particles/body) once a week using a MicroSprayer Aerosolizer and a high-pressure syringe (PennCentury, Philadelphia, PA, USA). The control mice were age-matched and exposed to air only. Emphysema and small airway remodeling were assessed at 6 months via morphometry on hematoxylin and eosin–stained and Masson’s trichrome–stained tissues, respiratory mechanics using the Flexivent system, and cell counts in bronchoalveolar lavage fluid samples. For the cigarette smoke and poly(I:C) exposure mouse models, 8-week-old mice were exposed to cigarette smoke for 21 days as described above. Additionally, mice received intranasal administrations of 50 μ poly(I:C) (Invivogen, tlrl-pic) on days 5, 9, 12, 16, and 19. Intratracheal administration of LF-EVs and LipoFB(R)-EVs (1.0×10^9^ particles/body) was conducted on days 7, 14, and 21. Equivalent volumes of PBS were used as controls. Age-matched air-exposed mice were included as non-smoking controls. Evaluation on the 23^rd^ day involved quantifying small airway remodeling through morphometry of hematoxylin and eosin–stained and Masson’s trichrome–stained tissue, respiratory mechanics, and cell counts in BAL fluid samples.

## PKH67-labeling of EVs

LF-derived EVs were labeled using a PKH67 green fluorescent labeling kit (Sigma-Aldrich, MINI67-1KT). The isolated EVs were incubated with 2 μM of PKH67 for 5 minutes and washed four times using a 100 kDa filter (Millipore, UFC510096) to remove excess dye and incubated with HBECs and AT2 cells for 24 hours. DAPI staining was carried out immediately before imaging. The EV uptake was evaluated using a fluorescence microscope (BZ-X810, Keyence).

## Hydroxyproline content of the whole lung

To quantitatively measure collagen in the left lungs of mice, a hydroxyproline assay was performed according to the manufacturer’s instructions (Chondrex, 6017). Left lung tissues from mice (10 mg) were homogenized in 100 μl of distilled water using PowerMasher II (Nippi, 891300) and BioMasherII (Nippi, 320103).

## Enzyme-linked immunosorbent assay (ELISA)

Mouse tissues were homogenized in ice-cold protein extraction reagent (Thermo Fisher Scientific, 78510) containing protease and phosphatase inhibitors. TNF-α concentration in lung tissue homogenate was measured using a mouse TNF-alpha Quantikine HS ELISA Kit (R&D Systems, MHSTA50), as per the manufacturer’s instructions.

## Amino acid assays

The total amount of free L-amino acids present in AT2 cells was determined using the L-amino acid assay kit (Cell Biolabs, MET-5054). The amino acid uptake activity of HBECs and AT2 cells was assessed using the amino acid uptake assay kit (DOJINDO, UP04). Amino acid uptake assay kit allows a simple fluorescence assay for amino acid transporter activity using Boronophenylalanine (BPA) and the fluorescent probe in this kit. These measurements were performed according to the manufacturer’s instructions. Absorbance and fluorescence were measured using a fluorescence microplate reader (Thermo Scientific, Varioskan LUX).

## [^3^H]L-leucine uptake and inhibition experiments

[^3^H]L-leucine uptake in the suspension cells was performed by rapid filtration with some modification^27,28^. HBECs cells (3.5 × 10^4^ cells) were washed twice with 37 °C pre-warmed Na^+^-free Hank’s balanced salt solution (HBSS) containing 125 mM choline-Cl, 25 mM HEPES, 4.8 mM KCl, 1.2 mM MgSO4, 1.2 mM KH2PO4, 1.3 mM CaCl2, and 5.6 mM glucose (pH 7.4). The uptake was measured for 1 minute at 37 °C in the same buffer containing 1 µM [^3^H]L-leucine (1 × 10^3^ Ci/mol). For inhibition experiments, 3 mM BCH was added to the radioisotope substrate. The uptake was terminated by adding ice-cold Na^+^-free HBSS, and the cells were filtrated through a 25 mm GF/B filter (Whatman, 1821-025), followed by washing with the same buffer. Membranes containing the cells were soaked in an Emulsifier-Safe cocktail (PerkinElmer). The radioactivity was measured with a β-scintillation counter (Hidex 600SL).

## Sample preparation for LC-MS

Proteins from 100 µL aliquots were dissolved in 100 µL of 2X Lysis Buffer (10 % SDS, 50 mM TEAB, pH 8.5). The total protein concentration was determined using the Micro BCA Protein Assay kit (Thermo Fisher Scientific). Sample preparation followed the protocol provided by the S-trap micro spin column (ProtiFi) with adaptations. Initially, samples were treated with 20 mM dithiothreitol, then boiled at 95 [for 10 minutes. After cooling to room temperature, samples were alkylated with 40 mM iodoacetamide (Thermo Fisher Scientific) and left to incubate at room temperature for 30 minutes in darkness. Subsequently, samples were acidified using phosphoric acid (FUJIFILM Wako Pure Chemical Corporation) and diluted with S-trap binding/wash buffer (100 mM TEAB, pH 7.1, Honeywell, in 90 % methanol). The samples were left at room temperature for 5 minutes. S-trap micro columns were prepared by equilibrating them with 150 µL of 0.2 % formic acid (FA) in 50 % acetonitrile, followed by centrifugation at 4000g for 1 minute, and then with 150 µL of S-trap binding/wash buffer, again followed by centrifugation. After equilibration, samples were transferred to the S-trap columns and centrifuged at 4000g for 1 minute. The protein amount added to each column was adjusted to 18.75 µg or less. The columns were washed four times with 200 µL of S-trap binding/wash buffer, with an additional 200 µL acetonitrile wash during the second wash. The trapped proteins were subjected to peptide digestion on the columns using 20 µL of Digestion Buffer (50 mM TEAB containing 750 ng of Trypsin/Lys-C Mix, Promega) at 47 [for 2 hours. Peptides were eluted with 40 µL of 50 mM TEAB, 40 µL of 0.2 % FA, and 35 µL of 0.2 % FA in 50 % acetonitrile. Each eluted fraction was combined in the same collection tube.

## Proteomic analysis with LC-MS

The reconstituted peptides (10 µL of water containing 0.1% formic acid) were quantified using the Pierce™ Quantitative Fluorometric Peptide Assay (Thermo Fisher Scientific). Samples were analyzed using an EASY-nLC 1200 System (Thermo Fisher Scientific) coupled to an Orbitrap Exploris 480 system (Thermo Fisher Scientific). Peptide samples (0.2 μg) were injected onto an Acclaim PepMap 100 trap column (75 μm × 2 cm, nanoViper C18 3 μm, 100Å, Thermo Fisher Scientific). The analytical column used was a C18 reverse-phase Aurora UHPLC Emitter Column with nanoZero (75 μm × 25 cm, Ion Opticks Pty Ltd) heated to 50 [using an ING ion source (AMR INCORPORATED). The nano pump flow rate was set to 300 nL/min with a 70-minute gradient using mobile phases A (0.1% FA in water) and B (95% acetonitrile and 5% water containing 0.1% FA). The chromatography gradient was designed as follows: 0-0.5 min at 3% B, 0.5-32.5 min from 3% B to 16% B, 32.5-45 min from 16% B to 24% B, 45-54 min from 24% B to 33% B, 54-55 min from 33% B to 95% B, followed by a 15-minute wash. Data-independent acquisition was conducted in positive ion mode. MS1 parameters included an m/z range of 500 to 860, resolution of 30,000, normalized AGC target of 300%, maximum injection time set to auto, and FAIMS CV voltage at -40. MS2 parameters included a resolution of 30,000, normalized AGC target of 300%, maximum injection time of 55 ms, collision energy of 30, isolation window of 6, with no overlapping (divided into 60, cycle time, 3.3 seconds), and FAIMS CV voltage at -40.

## DIA data analysis

All DIA samples were analyzed using DIA-NN (version 1.8) configured to search an in silico spectral library. The spectral library was generated by DIA-NN from the SwissProt Human database FASTA (downloaded October 2021). Parameters for creating the spectral library included an MS1 m/z range of 500-860, MS2 m/z range of 200-1800, fixed modification: carbamidomethyl(C), variable modifications: acetyl (protein N-term), oxidation(M), and M excision (protein N-term). Two missed trypsin cleavages and two modifications per peptide were allowed. Library search by DIA-NN was performed with parameters including Mass accuracy, MS1 accuracy, and Scan window set to 0, with MBR applied. The top six fragments were utilized for peptide identification and quantification. Protein quantification was calculated using the MAXLFQ algorithm. The FDR was set to 1% at both peptide precursor and protein group levels.

## Statistics and Data analysis

Data presented in bar graphs are shown as the average (±SEM) for technical replicates. Student’s t-test was used to compare the two datasets. An analysis of variance (ANOVA) test was used for multiple comparisons, followed by Tukey’s multiple comparison test to identify differences. Statistical significance was defined as *P* < 0.05. Prism version 9 (GraphPad Software, San Diego, CA, USA) was used for statistical analysis.

## Data availability

The protein expression profiles of human tissues were based on publicly accessible and downloadable data, available in the Human Protein Atlas (https://www.proteinatlas.org). The authors declare that the data supporting the findings of this study are available within the paper and its Supplementary Information file.

## Results

### A meta-analytical single-cell atlas of COPD lungs highlights metabolic dysregulation in LipoFBs

To elucidate the characteristics of LipoFBs in COPD, we first established a meta-analytical single-cell atlas of lung samples derived from both non-COPD patients and those with COPD. Following quality control of the scRNA-seq data, we integrated 12 datasets, including those from our previous research (**sTable1**)^5^. Primary LFs were clustered based on their expression profiles, and cell types were annotated based on established cell markers from the literature^21^. Employing Uniform Manifold Approximation and Projection (UMAP) clustering on 18,341 LFs from 160 samples revealed four distinct clusters (**Fig. 1A**). Among these clusters, we identified a subset characterized by LipoFBs that exhibited high expression of the genes, *Apolipoproteins E* (*APOE*) and *PLIN2* (**Fig. 1B**). Our exploration of the role of LipoFB in COPD lungs entailed a comparative analysis between COPD and non-COPD lungs (including non-COPD smokers and never-smokers). This investigation revealed no notable differences in the proportion of LipoFBs relative to the overall cell count or total fibroblast count between the two groups (**Fig. 1C**). Consequently, we embarked on a detailed examination of the differences in gene expression between LipoFBs isolated from individuals with and without COPD. IPA unveiled enrichment of the top 20 canonical pathways in LipoFBs derived from COPD lungs compared with those from non-COPD lungs. Within this set of pathways, seven were associated with glucose, lipid, and amino acid metabolism, including oxidative phosphorylation. Interestingly, most of these pathways exhibited dampened signals within LipoFBs derived from COPD samples. Furthermore, an additional group of seven signals related to protein synthesis displayed notable suppression within the COPD-derived LipoFBs across most involved pathways (**Fig. 1D**).

**Figure 1.**
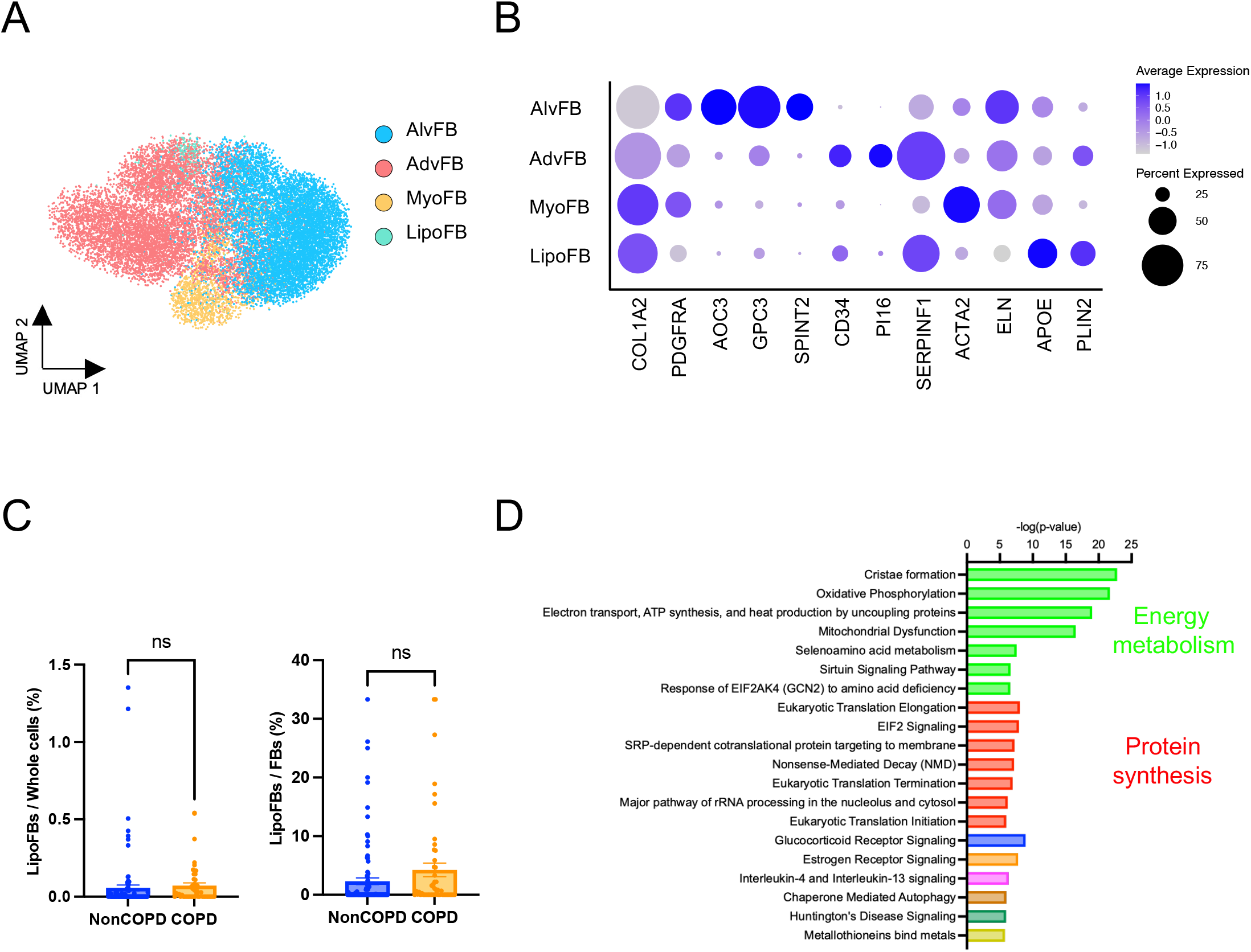
Lung fibroblast subtype clustering and characterization analyzed using meta-analytic single-cell RNA sequencing. (**A**) UMAP plot representing the clustering of LF subtypes based on SingleR annotation. Four distinct clusters were identified: alveolar fibroblast (AlvFB), adventitial fibroblast (AdvFB), myofibroblast (MyoFB), and lipofibroblast (LipoFB). (**B**) Dot plot showing the gene expression profiles of LF subtypes. Each dot represents the expression level of a gene within each subtype of LFs. (**C**) Graph comparing the percentage of LipoFB cells/whole cells or total LFs between non-COPD patients and those with COPD. (**D**) The top 20 canonical pathways derived from differential gene expression within the LipoFB cluster between non-COPD patients and those with COPD were analyzed using Ingenuity Pathway Analysis (IPA)

In healthy lungs, LipoFBs play a supportive role in AT2 cells in the alveolar microenvironment^9^. However, in the context of COPD pathogenesis, our analysis of the single-cell transcriptome suggested a potential disruption of this function in LipoFBs due to metabolic abnormalities. This finding implies a critical role for healthy LipoFBs in potentially preventing the onset and progression of COPD and being a therapeutic strategy by leveraging their secreted factors.

### Metabolic agents induce lipogenic differentiation in LFs

Given the exceedingly low proportion of LipoFBs in the human lungs, we endeavored to induce LipoFBs from LFs *in vitro*. Here, we aimed to circumvent the scarcity of LipoFBs within the pulmonary microenvironment, thereby facilitating a more comprehensive study and understanding of their characteristics and functions. Therefore, LFs from human lung specimens were isolated and cultured. Subsequently, induction of lipogenic differentiation with 5 mM Metformin or 100 μM Rosiglitazone was attempted (**Fig. 2A**). Metformin- or Rosiglitazone-treated LFs exhibited a distinctive morphology, characterized by the presence of droplets stored in their cytoplasm (**Fig. 2B**). The treatment of cells resulted in lipid droplet accumulation, as observed by fluorescence microscopy and flow cytometry and indicated by staining with the neutral lipid dye LipidTOX (**Fig. 2C, D**). Furthermore, qPCR analysis demonstrated a significant upregulation of the lipogenic marker genes, namely *Adipose differentiation-related protein* (*ADRP*), *CD36*, *Fibroblast growth factor 10* (*FGF10*), and *APOE* in LFs treated with Metformin or Rosiglitazone (**Fig. 2E**). In particular, the induction of *ADRP*, *CD36*, and *APOE* expression was notably higher in Rosiglitazone-treated LFs as compared to those treated with Metformin. Next, we explored potential variations in the induction of lipogenic differentiation across LF donor backgrounds: healthy controls, COPD, or IPF lungs. While minor differences were noted in lipid droplet accumulation (**sFig. 1A**) and lipogenic marker gene expression evaluation (**sFig. 1B**), the response patterns were notably consistent and significant across LFs, irrespective of their origin.

**Figure 2.**
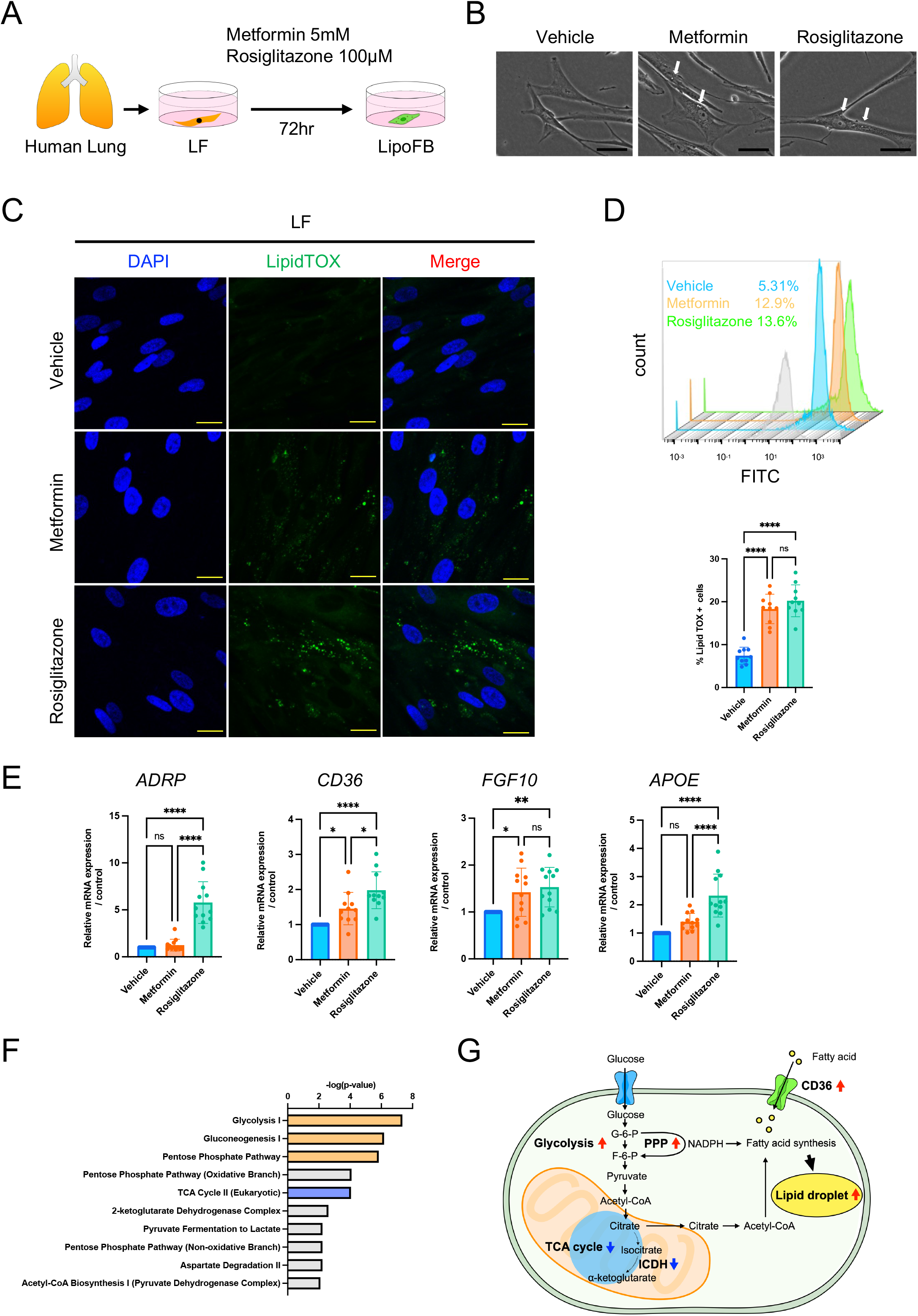
Induction and characterization of LipoFBs *in vitro* (A) Schematic representation of LipoFB induction by metabolic agents. (B) Representative phase contrast micrographs of LFs treated with Metformin, Rosiglitazone, or a vehicle. The white arrows represent lipid droplet-like structures. Scale bars = 50 μm. (C) Staining of lipid droplets in LFs treated with Metformin, Rosiglitazone, or the vehicle using LipidTOX (green). Nuclei were stained with DAPI (blue). Scale bars = 20 μm. (D) Representative flow cytometry histogram and quantification of LipidTOX^+^ cell abundance in response to Metformin or Rosiglitazone treatment. The gray histogram represents unstained control cells. \*\*\*\**P* <0.0001. ns; not significant. (E) qPCR analysis for the lipogenic marker genes, namely, *ADRP*, *CD36*, *FGF10*, and *APOE,* in LFs treated with Metformin, Rosiglitazone, or the vehicle. \*\*\*\**P* <0.0001, \*\**P* <0.01, \**P* <0.05. ns; not significant. (F) LC-MS performed on LFs treated with or without Rosiglitazone to identify proteins differentially expressed between LFs and LipoFBs. PEAKS and IPA software were used to analyze the proteomic data. (G) Schematic diagram of cellular metabolic changes in LipoFBs. G-6-P: glucose 6-phosphate, F-6-P: glucose 6-phosphate, PPP: pentose phosphate pathway, ICDH: Isocitrate Dehydrogenase, TCA: tricarboxylic acid cycle.

To elucidate the metabolic changes in LipoFBs, LC-MS analysis was conducted on LFs, both treated and untreated with Rosiglitazone (**sTable 2**). IPA analysis revealed an elevation in the glycolytic and pentose phosphate pathways (PPP), along with a decrease in the tricarboxylic acid (TCA) cycle within Rosiglitazone-induced LipoFBs (**Fig. 2F**). Upon further investigation, lowered levels of isocitrate dehydrogenase (ICDH) suggested the inhibition of the citric acid cycle. The addition of Rosiglitazone to LFs altered their metabolic pathway, shifting their pathways from ATP production via the citric acid cycle toward fatty acid synthesis and lipid droplet formation. Additionally, qPCR analysis indicated an increased expression of *CD36*, signifying the activation of fatty acid uptake (**Fig. 2E**). Collectively, these findings indicate that LipoFBs undergo metabolic reprogramming, wherein fatty acid synthesis is accelerated (**Fig. 2G**).

### LipoFB-derived EVs mitigate cigarette smoke-induced senescence and inflammation in lung epithelial cells

To investigate the impact of metabolic agents on lipogenic differentiation in LFs and the resulting changes in the secreted extracellular vesicles (EVs), LipoFB-derived EVs (LipoFB-EVs) and LF-derived EVs (LF-EVs) were isolated by conventional ultracentrifugation of conditioned medium collected from these cells and extensively characterized (**sFig. 2A**). The particle size and quantities of the EV preparations measured by nanoparticle-tracking analysis showed no significant differences between LipoFB-EVs and LF-EVs (**sFig. 2B, C**). In addition, cryo-transmission electron microscopy revealed typical bilayer membrane vesicles with a size range of 50-150 nm in both sets of EV isolates (**sFig. 2D**). Western blot analysis revealed the presence of the EV marker proteins CD9, CD63, and Caveolin-1 in both sets of EVs, whereas the actin cytoskeleton and mitochondrial protein TOM20 were not enriched in the EVs (**sFig. 2E**). These EVs fulfilled the minimal experimental criteria for EV identification and were mainly composed of small EVs, as described in the position statement of the International Society for Extracellular Vesicles (MISEV2023)^30^.

To delineate the antiaging attributes of these EVs during COPD treatment^31^, we assessed their capacity to mitigate cellular senescence in lung epithelial cells. Cigarette smoke (CS) is known to induce lung epithelial cell senescence, potentially contributing to the pathogenesis of airway remodeling in COPD. In an experimental setting, exposure to low concentrations of cigarette smoke extract (CSE) induces cellular senescence in human bronchial epithelial cells (HBECs). Although we found that LF-EVs and Metformin-induced LipoFB (LipoFB(M))-EVs had no remarkable effect on the expression of the senescence markers p21 and p16 in HBECs exposed to 1.0% CSE, Rosiglitazone-induced LipoFB (LipoFB(R))-EVs clearly inhibited the CSE-mediated induction of p21 and p16 expression (**Fig. 3A**). In addition, CSE-induced expression of senescence-associated-β-galactosidase (SA-β-gal) staining was also suppressed only by LipoFB(R)-EVs (**Fig. 3B**). The DNA damage response plays a critical role in initiating cellular senescence and is activated through the p53/p21 and/or p16/pRb pathways^32^. In LipoFB(R)-EVs, the reduction in the count of γ-H2AX (phosphorylated Histone H2A.X)-positive nuclei indicates the suppression of the DNA damage response (**sFig. 3A**). To further delineate the mechanisms influenced by LipoFB-EVs that contribute to the attenuation of senescence in CSE-treated HBECs, we assessed ROS production within the cells. Notably, CSE-exposed HBECs incubated with LipoFB(R)-EVs exhibited reduced intracellular ROS levels, which was quantified using the 2’,7’-dichlorofluorescin diacetate (DCFH-DA) assay (**Fig. 3C**). These findings suggest the involvement of ROS in the mechanisms responsible for mitigating the DNA damage response and slowing senescence in CSE-exposed HBECs, mediated by LipoFB(R)-EVs.

**Figure 3.**
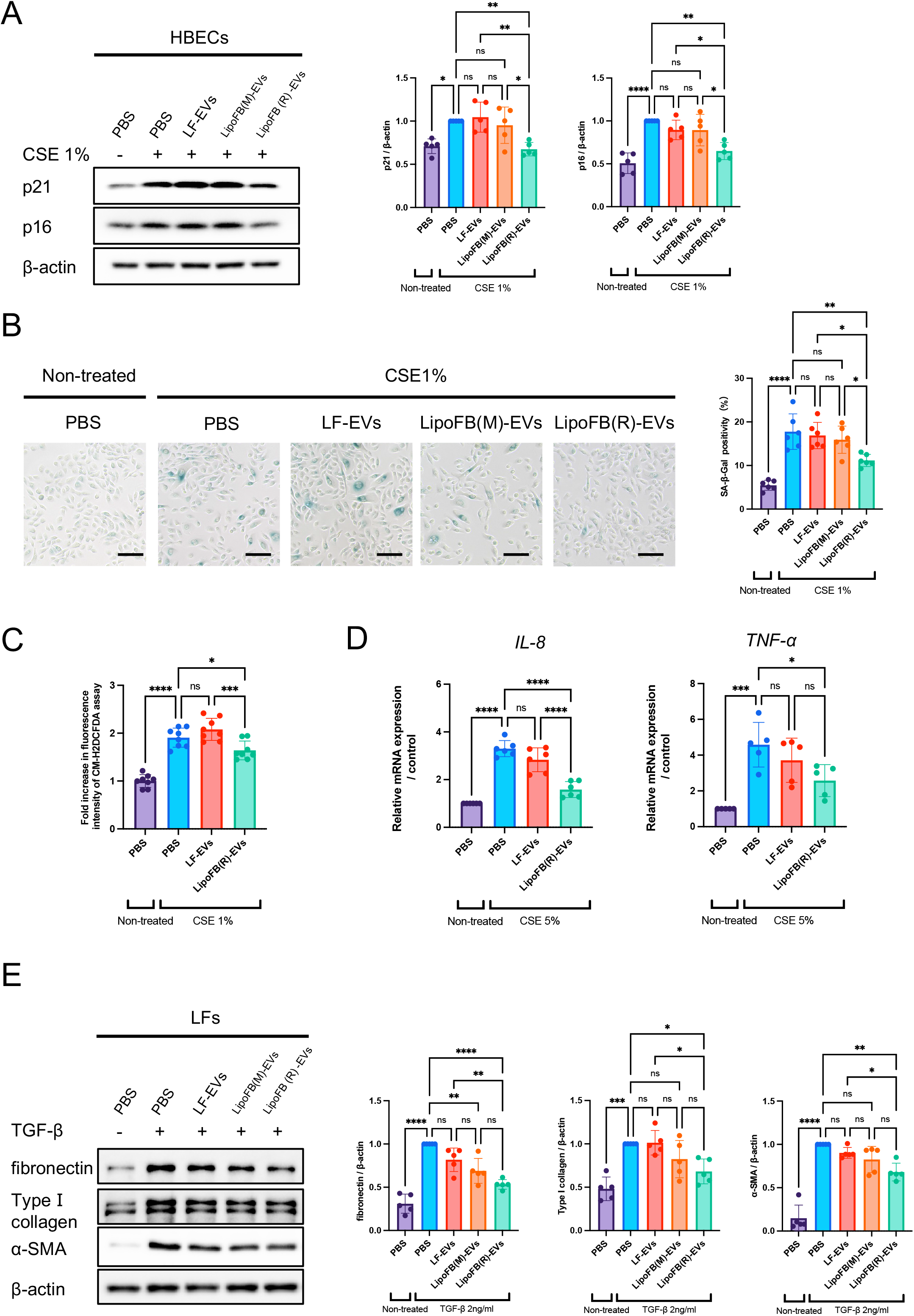
LipoFB-EVs attenuate CS and TGF-β-induced COPD pathologies *in vitro* (A) Representative immunoblot and quantitative analysis showing the amount of p21, p16, and β-actin in HBECs treated for 48 hours with PBS, LF-EVs, LipoFB(M)-EVs, or LipoFB(R)-EVs (10 μg/ml) in the presence or absence of CSE (1.0%). \*\*\*\**P* <0.0001, \*\**P* <0.01, \**P* <0.05. ns; not significant. (B) Photomicrographs and quantitative analysis of SA-β -gal staining of HBECs treated for 48 hours with PBS, LF-EVs, LipoFB(M)-EVs, or LipoFB(R)-EVs (10 μg/ml) in the presence or absence of CSE (1.0%). \*\*\*\**P* <0.0001, \*\**P* <0.01, \**P* <0.05. ns; not significant. Scale bars = 100 μm. (C) Fluorescence intensity of DCFH-DA staining for intracellular ROS production of HBECs in response to PBS, LF-EVs, or g/ml) in the presence or absence of CSE (1% for 48 hours). \*\*\*\**P* <0.0001, \*\*\**P* <0.001, \**P* <0.05. ns; not significant. (D) Measurement of cytokine secretion in HBECs in response to PBS, LF-EVs, or LipoFB(R)-EVs (10 μg/ml) in the presence or absence of CSE (5% for 8 hours). \*\*\*\**P* <0.0001, \*\*\**P* <0.001, \**P* <0.05. ns; not significant. (E) Representative immunoblot and quantitative analysis showing the amount of fibronectin, type I collagen, α-SMA, and β-actin in LFs treated for 24 hours with PBS, LF-EVs, LipoFB(M)-EVs, or LipoFB(R)-EVs (10 g/ml) in the presence or absence of TGF- (2 ng/ml for 24 hours). \*\*\*\**P* <0.0001, \*\*\**P* <0.001, \*\**P* <0.01, \**P* <0.05. ns; not significant.

Another pathology induced by CS exposure involves an aberrant inflammatory response, which is a key component in the pathogenesis of COPD^33^. In experimental conditions, exposure to a low concentration of CSE-induced inflammation in HBECs. While LF-EVs demonstrated no significant impact on IL-8 and TNF-α expression in HBECs exposed to 5.0% CSE, LipoFB(R)-EVs distinctly inhibited the aberrant inflammatory response in these cells (**Fig. 3D**).

### LipoFB-derived EVs attenuate TGF-β -induced myofibroblast differentiation

Airway remodeling, a result of inflammation and fibrosis, is a significant aspect of COPD pathology. To characterize the antifibrotic properties of these EVs in COPD treatment, we utilized LFs to assess their impact on TGF-β -induced myofibroblast differentiation. Indicated by the increased expression of fibronectin, type I collagen, and α-SMA, LipoFB(R)-EVs significantly countered TGF-β-induced myofibroblast differentiation; this response is not seen with LF-EVs or LipoFB(M)-EVs (**Fig. 3E**). Moreover, LipoFB(R)-EVs inhibited myofibroblast differentiation in a dose-dependent manner (**sFig. 3B**). Subsequently, we compared the inhibitory potency of LipoFB(R)-EVs against pirfenidone (PFD) and nintedanib (NTD). The approximate maximum drug concentrations (Cmax) in clinical use are 10 μg/ml for PFD and 60 nM for NTD. In comparison to doses of 10 μg/ml PFD and 100 nM NTD, LipoFB(R)-EVs demonstrated a more potent suppression of TGF-β-induced myofibroblast differentiation in LFs. To clarify the specific activity of LipoFB(R)-EVs in suppressing myofibroblast differentiation, we examined the antifibrotic properties of EVs derived from LFs treated with various metabolic agents. Comparing the effects of EVs derived from LFs treated with not only Metformin but also Pioglitazone, Ciglitazone, Dibutyryl cyclic AMP (DBcAMP), Parathyroid hormone-related peptide (PTHrP), Pemafibrate, or Pravastatin, it was evident that LipoFB(R)-EVs exhibited the most potent antifibrotic effects on myofibroblast differentiation (**sFig. 3C**). Taken together, this data suggests that EVs secreted by Rosiglitazone-induced metabolically reprogrammed LFs demonstrate superior antifibrotic effects and show immense potential as promising candidates for COPD therapeutics.

### LipoFB-derived EVs promote AT2 stem cell restoration

The impairment of AT2 cells is a pivotal step in the pathogenesis of COPD. To assess the role of AT2 cells in COPD lungs, we isolated AT2 cells derived from human lung samples (**sFig. 4A**) and employed AT2 organoids in feeder-free 3D cultures^26^. Indeed, the colony-forming efficiency (CFE) of AT2 organoids derived from the lungs of patients with COPD was significantly lower than that of those derived from healthy lungs (**Fig. 4A**). Furthermore, we examined the effects of LipoFB(R)-EVs on AT2 organoids. We confirmed the uptake of lipophilic dye PKH67-labeled EVs into AT2 organoid cells (**Fig. 4B**). Notably, LipoFB(R)-EVs increased the size of AT2 organoids and bolstered the viability of AT2 cells (**Fig. 4C**). Similar effects were observed in both 2D-cultured HBECs (**sFig. 4B**) and 3D-cultured HBEC organoids (**sFig. 4C, D**). The HBECs established in our protocol expressed basal cell markers such as transformation-related protein 63 (Trp63) and cytokeratin 5 (Krt5)^34^, suggesting that the effects of LipoFB(R)-EVs might extend beyond AT2 cells, potentially impacting a broader spectrum of epithelial stem cells.

**Figure 4.**
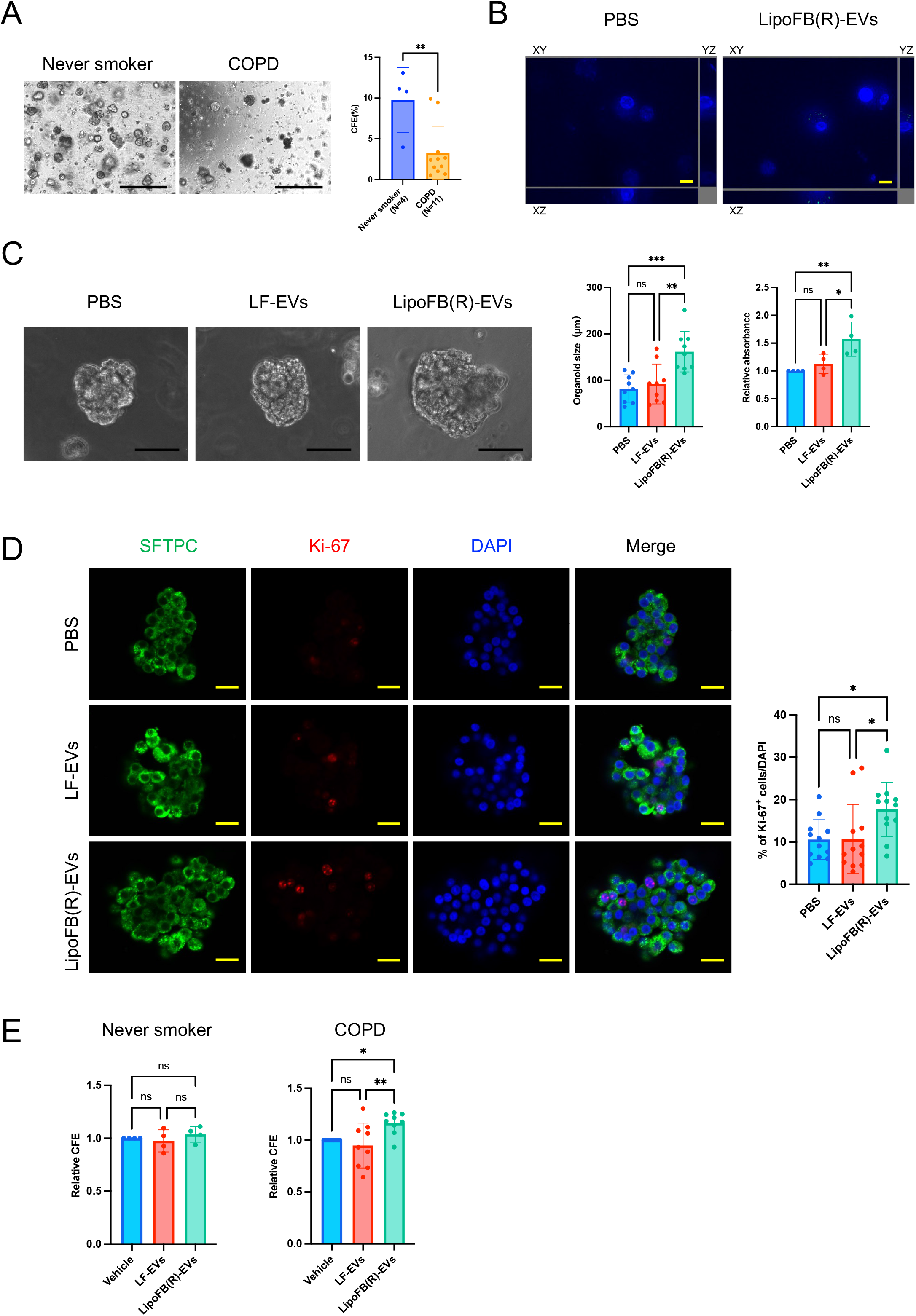
LipoFB-EVs can restore AT2 stemness (A) Representative images and graphs of the colony formation efficiency (CFE) of AT2 cells from healthy subjects (n=4) and patients with COPD (n=11) respectively. Scale bars = 500 μm. \*\**P* <0.01. (B) Confocal microscopy z-stack images of AT2 organoids incubated with PKH67[labeled LipoFB(R)-EVs (green). z-stack section (x, y) at the level of AT2 cells with orthogonal x (y, z) and y (x, z) sections on the right and bottom, respectively. Scale bars = 10 μm. (C) Cell viability of AT2 cells in 3D organoid cultures. Representative images, cell counting kit 8 (CCK-8) assay, and organoid size of AT2 organoids treated with PBS, LF-EVs, LipoFB(R)-EVs. \*\*\**P* <0.001, \*\**P* <0.01, \**P* <0.05. ns; not significant. Scale bars = 50 μm. (D) Whole mount staining and confocal imaging showing localization of SFTPC and Ki-67 in 3D cultures of AT2 organoids treated with PBS, LF-EVs, LipoFB(R)-EVs, and their quantitative graphs. \**P* <0.05. ns; not significant. Scale bars = 20 μm. (E) CFE of AT2 organoids treated with PBS, LF-EVs, and LipoFB(R)-EVs in healthy subjects and patients with COPD, respectively. **P <0.01, \**P* <0.05. ns; not significant.

Subsequently, immunofluorescence staining for SFTPC and Ki-67 was performed on AT2 organoids. The proliferation of SFTPC^+^ AT2 cells, as identified by Ki-67 expression, was significantly increased in LipoFB(R)-EV-treated AT2 organoids as compared to those treated with LF-EVs or the control (**Fig. 4D**). Importantly, these observations were consistent across AT2 organoids, indicating a consistent effect irrespective of their origin. However, a difference based on the lung background was observed only in CFE. Intriguingly, while no significant difference was observed in CFE in AT2 organoids derived from non-smoker lungs following LipoFB(R)-EV treatment, a significant enhancement in CFE was noted in AT2 organoids derived from COPD lungs following LipoFB(R)-EV treatment (**Fig. 4E**). This suggests that LipoFB(R)-EVs potentially augment the stemness of impaired AT2 cells in COPD lungs, thereby promoting colony formation by a single AT2 cell. These findings indicate that LipoFB(R)-EVs have the potential to enhance the viability and colony-forming ability of AT2 organoid models, suggesting their ability to promote AT2 stem cell restoration.

### LAT1 encapsulated within LipoFB-EVs demonstrates therapeutic potential for COPD traits in vitro

To unravel the underlying mechanisms, we conducted a proteomic analysis of LipoFB(R)-EVs and LF-EVs using liquid chromatography-mass spectrometry (LC-MS). The analysis identified 2,009 proteins encompassing markers typical of exosomes/small EVs, such as CD9, CD63, CD81, and TSG101 (**Fig. 5A**). Among these proteins, 1,025 exhibited significantly increased expression, while 984 proteins showed decreased expression in LipoFB(R)-EVs as compared to LF-EVs (**sTable 2**). Notably, the volcano plot revealed statistically significant differences, with 46 proteins displaying increased expression (fold change (FC) ≥1.5) and 19 exhibiting decreased expression (FC ≤ LipoFB(R)-EVs, when compared to LF-EVs (*P*<0.05) (**Fig. 5A, sFig. 5A**). Noteworthy among the 46 proteins with markedly elevated expression in LipoFB(R)-EVs are 6 proteins (L-type amino acid transporter 1 (LAT1)^35,36^, Hepatocyte growth factor (HGF)^37^, Niemann-Pick type C1 (NPC1)^38^, Thioredoxin reductase 1 (TrxR1)^39^, Angiopoietin-like 4 (ANGPTL4)^40^, and Collagen triple helix repeat containing protein 1 (CTHRC1)^41^) previously implicated in the functionality of AT2 cells (**Fig. 5A, sFig. 5A**). Among the six identified proteins, we focused on LAT1 (SLC7A5), which plays a pivotal role in facilitating the transport of essential neutral amino acids, including leucine, isoleucine, valine, phenylalanine, tyrosine, tryptophan, methionine, and histidine. Building upon previous findings, PPARγ has been reported to induce LAT1 through the mediation of specificity protein 1(SP-1)^42^. Furthermore, the effective localization of LAT1 on the cell membrane necessitated its interaction with 4F2hc (SLC3A2), which was also encompassed within this set of 46 proteins (**Fig. 5A, sFig. 5A**). Amino acid transporters, such as LAT1, are pivotal for cell growth and operate within a sodium-independent large neutral amino acid transport system^43^. Immunoblotting analysis confirmed higher levels of LAT1 and 4F2hc both in cells and EVs derived from LipoFB(R) than in LFs (**Fig. 5B, sFig. 5B**). Furthermore, following the treatment of AT2 organoids with LipoFB(R)-EVs, a significantly greater increase in LAT1 and 4F2hc expression was observed in AT2 organoids than in those treated with LF-EVs (**Fig. 5C, sFig. 5C**). Under non-reducing conditions (without DTT), both antibodies against LAT1 and 4F2hc identified approximately a 125-kDa protein (**sFig. 5D**). Conversely, under reducing conditions (with DTT), approximately a 37-kDa protein for LAT1 and 75-/100-kDa proteins for 4F2hc were observed. This confirms that AT2 organoids treated with LipoFB(R)-EVs highly expressed both LAT1 and 4F2hc, forming a heterodimer *via* a disulfide bond (**Fig. 5C, sFig. 5D**). While exploring the potential influence of EV-mediated regulation on LAT1 transcriptional levels, including miRNA effects, qRT-PCR analysis revealed no observable changes at 6, 12, or 24 hours after EV treatment (**sFig. 5E**). This observation suggests the transfer of the LAT1 protein by EVs to AT2 cells. Recent studies have indicatedb disruptions in amino acid homeostasis in patients with COPD, showing changes in various amino acid levels in the blood and tissues^44^. Specifically, alterations in amino acid profiles, including decreases in branched-chain amino acids (BCAAs) such as leucine, isoleucine, and valine, have been observed^45^. Our examination of the total amino acid content in AT2 cells sourced from both COPD and non-COPD (including non-COPD smokers and never-smokers) lungs revealed a significant reduction, specifically within AT2 cells derived from COPD lungs (**Fig. 5D**). Indeed, the gene expression of LAT1, obtained from the meta-analytic single-cell atlas of lung samples derived from both COPD and non-COPD lungs (**sTable1**), was significantly reduced in the AT2 fraction of lungs from patients with COPD as compared to that in lungs from non-COPD patients (**Fig. 5E**). Furthermore, we confirmed that pathway analysis using meta-analytic single-cell RNA sequencing data showed a significant decrease in the expression of the metabolism of amino acids and derivatives in AT2 cells of patients with COPD as compared to non-COPD AT2 cells (**Fig. 5F, sTable 4**).

**Figure 5.**
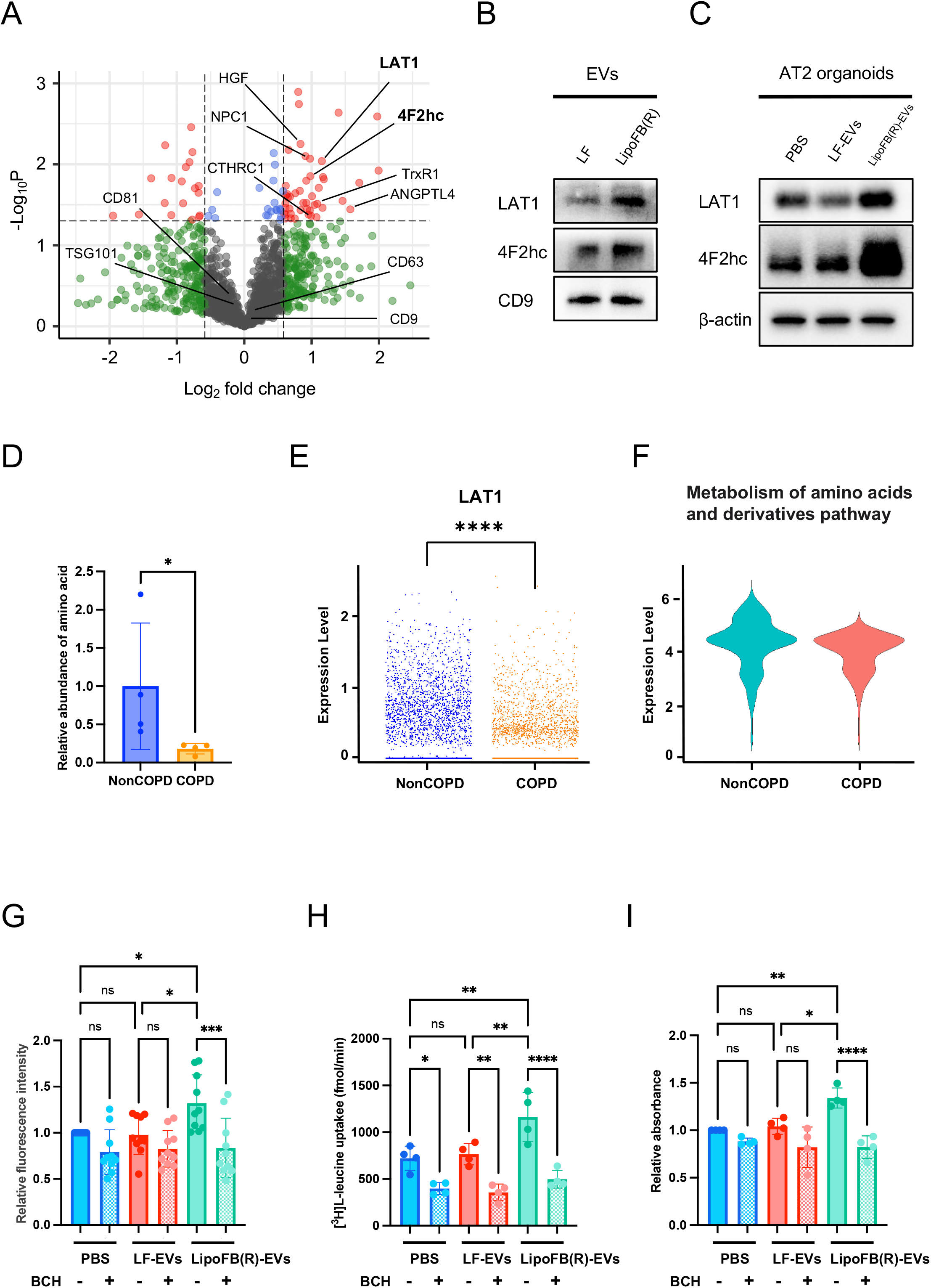
LAT1 encapsulated within LipoFB-EVs demonstrates a therapeutic potential for the impairment of AT2 cell stemness in COPD lungs (**A**) Volcano plot of statistically significantly different proteins identified by LC-MS between LF-EVs and LipoFB(R)-EVs. In the volcano plot, red dots represent proteins with a significant fold change (FC) ≥ 1.5 or≤ 0.667 and *P* < 0.05; blue dots represent proteins with a significant FC < 1.5 or > 0.667 and *P* < 0.05; green dots represent proteins with a significant FC ≥ 1.5 or ≤ 0.667 and *P* ≥ 0.05; black dots, no obvious changes in proteins. (**B**) Representative immunoblot of LAT1, 4F2hc, and CD9 in LF-EVs and LipoFB(R)-EVs. (**C**) Representative immunoblot of LAT1, 4F2hc, and β-actin in AT2s cell lysate treated with PBS, LF-EVs, or LipoFB(R)-EVs. (**D**) Comparison of amino acid content in AT2 cells isolated from the lungs of non-COPD (n=4) and COPD (n=4) patients respectively. \**P* <0.05. (**E**) The differential expression of the LAT1 gene in the AT2 cell population between non-COPD patients and those with COPD, analyzed using the meta-analytical single-cell RNA sequencing data. \*\*\*\**P* <0.0001. (**F**) Comparison of the expression of the metabolism of amino acids and their derivatives pathway from Reactome Pathway Database (https://reactome.org) in AT2 in non-COPD patients and those with COPD using the meta-analyticalsingle-cell RNA sequencing data. (**G**) Relative fluorescence intensity of amino acid uptake assays with or without BCH (3 mM) in AT2 cells treated with PBS, LF-EVs, or LipoFB®-EVs under sodium-free conditions. \*\*\**P* <0.001, \**P* <0.05. ns; not significant. (**H**) [^3^H]L-leucine uptake experiment in the presence or absence of BCH (3 mM) in AT2 cells treated with PBS, LF-EVs, or LipoFB(R)-EVs under sodium-free conditions. \*\*\*\**P* <0.0001, \*\**P* <0.01, \**P* <0.05. ns; not significant. (**I**) Inhibition assay using BCH (3 mM) for cell viability of AT2 cells. Cell counting kit 8 (CCK-8) assays of AT2 organoids treated with PBS, LF-EVs, or LipoFB(R)-EVs. \*\*\*\**P* <0.0001, \*\**P* <0.01, \**P* <0.05. ns; not significant.

To assess the involvement of LAT1 in the mechanism of action of LipoFB(R)-EVs as a potential COPD therapeutic agent, we employed 2-Amino-2-norbornanecarboxylic acid (BCH) under sodium-free conditions as an inhibitor of system L, including LAT1 and LAT2. In general, LAT1 is the only expressed transporter in the lungs, whereas LAT2 expression is absent (**sFig. 5F**). In the present study, the specific inhibitory effect of BCH under sodium-free conditions was attributed to LAT1. Importantly, amino acid uptake assays, using p-Boronophenylalanine (BPA) as a substrate^46^, revealed substantial values facilitated by LipoFB(R)-EVs added to AT2 organoids, and the uptake was diminished upon the addition of BCH (**Fig. 5G**). Furthermore, ^3^H-labeled leucine uptake experiments showed notable leucine uptake, which was also reduced upon the addition of BCH to in AT2 organoids (**Fig. 5H**). In AT2 organoid viability assessments, LipoFB(R)-EVs notably enhanced cell viability, a response that was confirmed to be nullified upon BCH addition (**Fig. 5I**). Moreover, amino acid uptake mediated by EV LAT1 transfer was observed in AT2 cells and other cell types, such as HBECs (**sFig. 5G, H**) and LFs (**sFig. 5I**). In HBEC organoid viability assessments, LipoFB(R)-EVs notably enhanced cell viability, a response that was confirmed to be nullified upon BCH addition (**sFig. 5J**).

Taken together, these findings suggest that the functionality of LAT1 encapsulated within LipoFB(R)-EVs in recipient cells demonstrates therapeutic efficacy through amino acid uptake, particularly by targeting the impaired AT2 cell stemness in COPD lungs.

### LipoFB-EVs block the airway pathology in a COPD model exacerbated by CS exposure

We assessed the therapeutic potential of LipoFB(R)-EVs in two different experimental mouse models of COPD. First, we established a murine model of COPD by combining short-term CS exposure with the intranasal administration of viral mimetic poly(I:C). The three-week regimen of CS exposure coupled with poly(I:C) treatment (**sFig. 6A**) resulted in increased airway inflammation and remodeling. Treatment with LipoFB(R)-EVs elicited substantial alterations in body weight as compared to both LF-EVs treatment and PBS control groups (**sFig. 6B**). We observed a significant increase in the total bronchoalveolar lavage (BAL) cell count, specifically that of macrophages, neutrophils, and lymphocytes in the BAL, alongside escalated expression of the lung inflammatory mediator TNF-α, and a decreased percentage of forced expiratory volume in 0.1 Ssecond (FEV0.1%: FEV0.1/FVC) (indicative of airway obstruction) in the CS-induced exacerbation model of COPD (**sFig. 6C-E**). The staining observations highlighted that the combined CS-poly(I:C) exposure amplified inflammation and led to small airway fibrosis in the murine COPD model (**sFig. 6F**), thus replicating the pivotal aspects of airway pathology observed in human COPD. During the regimen, an intratracheal injection of 1 × 10^9^ EV was administered on days 7, 14, and 21. Notably, the intratracheal administration of LipoFB(R)-EVs (but not LF-EVs) inhibited these exacerbations (**sFig. 6C-F**). Consistent with *in vitro* experiments, these results underscore the ability of LipoFB(R)-EVs to alleviate airway remodeling, inflammation, and obstructive airway physiology in a model simulating COPD exacerbation due to CS.

**Figure 6.**
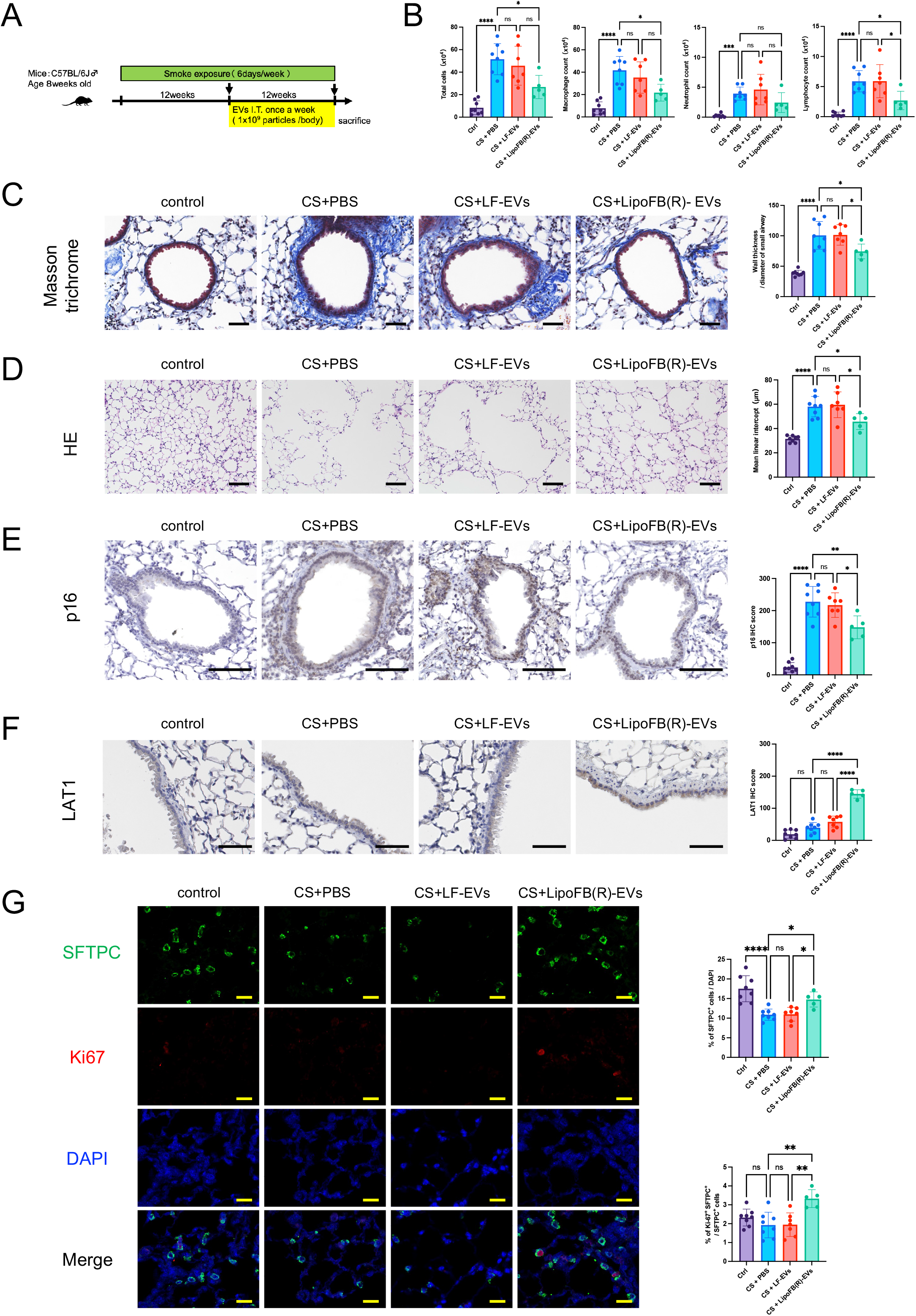
Intratracheal administration of LipoFB-EVs attenuates long-term cigarette smoke-induced COPD pathologies in mice (**A**) Schematic protocol for intratracheal (I.T.) EV treatment in the mouse model of COPD induced by 6 months of cigarette smoke (CS). (**B**) Cell counts of total cells, macrophages, neutrophils, and lymphocytes in the bronchoalveolar lavage fluid (BALF) from mice. n=8 in the control group, n=8 in the CS+PBS group, n=7 in the CS+LF-EVs group, and n=5 in the CS+LipoFB(R)-EVs group. (**C**) Representative images of Masson’s trichrome staining of lung tissue sections from each indicated group of treated mice. Quantification of the ratio of small airways wall thickness to the diameter of small airways. \*\*\*\**P* <0.0001, \**P* <0.05. ns; not significant. Scale bars = 50 μm. n=8 in the control group, n=8 in the CS+PBS group, n=7 in the CS+LF-EVs group, and n=5 in the CS+LipoFB(R)-EVs group. (**D**) Representative images of H&E staining of lung tissue sections from each indicated group of treated mice. Quantification of mean linear intercept (MLI). \*\*\*\**P* <0.0001, \**P* <0.05. ns; not significant. Scale bars = 100 μm. n=8 in the control group, n=8 in the CS+PBS group, n=7 in the CS+LF-EVs group, and n=5 in the CS+LipoFB(R)-EVs group. (**E**) Immunohistochemical staining of p16 in representative lung sections from each indicated group of treated mice. Quantification of IHC scores. Scale bars = 100 μ CS+PBS group, n=7 in the CS+LF-EVs group, and n=5 in the CS+LipoFB(R)-EVs group. (**F**) Immunohistochemical staining of LAT1 in representative lung sections from each indicated group of treated mice. Quantification of the percentage of DAB-positive cells. Scale bars = 50 μm. n=8 in the control group, n=8 in the CS+PBS group, n=7 in the CS+LF-EVs group, and n=5 in the CS+LipoFB(R)-EVs group. (**G**) Representative images of immunofluorescence co-staining for SFTPC (green) and Ki-67 (red) and the lung sections counterstained with DAPI (blue). Scale bars = 20 μm. Quantification of the number of AT2 cells and Ki-67+/SFTPC+ dual positive cells. \*\*\*\**P* <0.0001, \*\**P* <0.01, \**P* <0.05. ns; not significant. n=8 in the control group, n=8 in the CS+PBS group, n=7 in the CS+LF-EVs group, and n=5 in the CS+LipoFB(R)-EVs group.

### LipoFB-EVs attenuate emphysematous changes in a long-term CS exposure mouse model

Next, we developed a murine model of COPD through extended exposure to CS for more than 6 months (**Fig. 6A**). Control mice were exposed to ambient air. Long-term CS exposure resulted in increased airway remodeling, obstruction, accelerated cellular senescence, and alveolar emphysema in the model. This manifested as a notable increase in the total BAL cell count, particularly lymphocytes, accompanied by increased airway resistance (Rn) and elastance (Ers), reduced dynamic compliance (Crs), and a decline in FEV0.1%, indicating established airway obstruction due to prolonged CS exposure in our murine model (**Fig. 6B, sFig. 7A**). Histological assessments revealed intensified lung epithelial cellular senescence, as assessed by p16 expression levels, airway wall remodeling, and the induction of alveolar emphysema in the mean linear intercept within the murine COPD model (**Fig. 6C, D, E**). These findings closely mirror the significant aspects of airway and alveolar pathology observed in human COPD. For the last three months of the six-month duration, we administered intratracheal injections of 1 × 10^9^ EV particles once a week. Notably, intratracheal administration of LipoFB(R)-EVs (but not LF-EVs) mitigated these exacerbations (**Fig. 6B-E**). However, no significant differences in respiratory function parameters or body weight were observed between treatment interventions in each mouse group. (**sFig. 7A, B**). To validate the therapeutic effects of LipoFB(R)-EVs observed *in vitro*, LAT1 expression in the airways and alveoli of mice was evaluated. Although LAT1 expression was not observed in the lung parenchyma of control mice, control CS-treated mice, or LF-EV-treated mice, LipoFB(R)-EVs significantly increased the number of LAT1^+^ lung epithelial cells (**Fig. 6F**). To further identify the regenerative therapeutic effects of LipoFB(R)-EVs on alveolar emphysema, mouse lungs were stained via immunofluorescence techniques for SFTPC and Ki-67.

**Figure 7.**
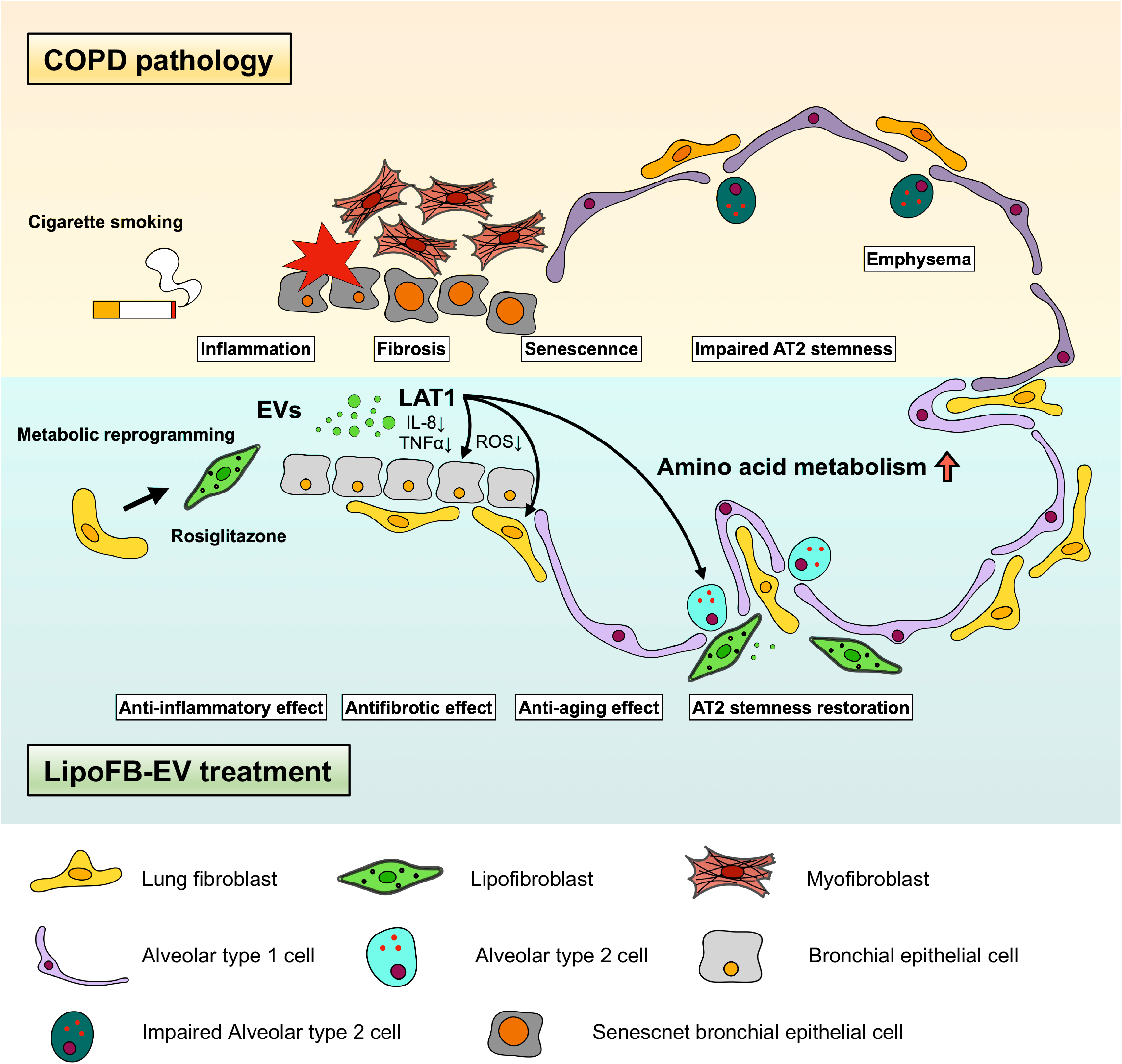
Schematic diagram of the proposed mechanism of the therapeutic potential of LipoFB-EVs against COPD pathologies LipoFB-EVs are taken up by AT2 cells, airway epithelial cells, and fibroblasts. They can improve COPD pathologies by transporting exosomal LAT1, thereby increasing cellular uptake of amino acids.

Crucially, while a notable decrease in the proportion of SFTPC^+^ AT2 cells was observed in control CS-treated mice, treatment with LipoFB(R)-EVs in CS-exposed mice significantly increased the proportion of SFTPC^+^ AT2 cells. Additionally, the proliferation of SFTPC^+^ AT2 cells, identified by Ki-67 expression, was significantly higher in LipoFB(R)-EV-treated mice than in LF-EV-treated or control CS-exposed mice (**Fig. 6G**). Accordingly, these results suggest the therapeutic potential of LipoFB(R)-EVs in alleviating airway remodeling, accelerated cellular senescence, and alveolar emphysema through LAT1 transfer in a murine model of COPD induced by long-term CS exposure.

## Discussion

This study highlighted the promising role of LipoFB-EVs containing LAT1 in the treatment of COPD. These LipoFB-EVs, particularly those generated by Rosiglitazone treatment, have diverse therapeutic potentials. They effectively mitigated key COPD characteristics, such as cellular senescence and inflammatory responses, in lung epithelial cells induced by CSE. This was achieved by reducing ROS and modulating DNA damage response pathways, suggesting that LipoFB-EVs are a potential treatment for COPD. Moreover, LipoFB-EVs demonstrate antifibrotic properties by inhibiting TGF-β-induced myofibroblast differentiation, surpassing conventional antifibrotic drugs. They also aid in restoring impaired AT2 stem cells, which are crucial for lung homeostasis, by enhancing their viability, colony-forming ability, and proliferation. Upon administration via intratracheal injection in murine COPD models, LipoFB-EVs alleviated airway inflammation, remodeling, obstruction, cellular senescence, and alveolar emphysema induced by both short- and long-term CS exposure. This therapeutic efficacy is attributed to the transfer of LAT1, a pivotal amino acid transporter, via LipoFB-EVs to recipient cells. LAT1 facilitates cell growth, proliferation, and stemness maintenance through amino acid uptake mechanisms, particularly leucine transport, thereby aiding in the restoration of lung function in patients with COPD. Overall, encapsulating LAT1 within LipoFB-EVs has emerged as a novel therapeutic strategy for COPD, targeting various pathological mechanisms, such as cellular senescence, inflammation, fibrosis, and stem cell dysfunction, offering a comprehensive approach to COPD management (**Fig. 7**).

In the present study, we explored the role of LipoFBs in COPD pathogenesis. Transcriptomic analysis revealed metabolic irregularities in LipoFBs in the lungs of patients with COPD Previous reports suggested that LipoFBs transfer neutral lipids to AT2 cells, aiding in surfactant phospholipid synthesis via ADRP. This process supports AT2 proliferation and their differentiation into AT1 cells to maintain alveolar homeostasis^47,48^. Our investigation demonstrates the therapeutic potential of EVs derived from Rosiglitazone-induced LipoFBs in COPD pathogenesis, including AT2 stem cell restoration. Although further investigation is required, these findings suggest that metabolic irregularities in LipoFBs may disrupt AT2 stemness through the transfer of neutral lipids and EVs, thereby potentially influencing the initiation and progression of COPD.

The PPARγ agonist Rosiglitazone, a thiazolidinedione-class medication, is commonly employed for managing type 2 diabetes by enhancing insulin sensitivity. In addition, addressing lipid abnormalities has been shown to be effective. However, concerns have been raised regarding its association with elevated severe side effects, particularly cardiovascular risks, necessitating caution and close monitoring during its use. The half-maximal effective concentration (EC50) of rosiglitazone reported so far is 11 nM^49^. Despite the notably elevated concentrations used *in vitro*, including those in this study, achieving high-dose administration for lipid differentiation in LFs remains a challenge. Therefore, we have devised a strategic approach in which elevated concentrations of Rosiglitazone were applied i*n vitro* to induce metabolic reprogramming in LFs. This process culminated in the collection of LipoFB-EVs for use in novel therapeutic approaches. This innovative therapeutic strategy, which utilizes the *in vitro* high-dose administration of Rosiglitazone solely for cell engineering to obtain therapeutic EVs, represents a novel and potentially versatile approach for drug repositioning. This method holds promise as a generalizable approach for novel therapeutic applications as it overcomes the challenge of administering high doses of various metabolic regulators to humans.

In this study, we have elucidated the diverse therapeutic effects of LipoFB-EVs on COPD. Elevated levels of LAT1-4F2hc heterodimer in LipoFB-EVs may contribute to amino acid uptake and cellular effects in the recipient cells. Specifically, LipoFB-EVs demonstrated the ability to mitigate smoking-induced aging and inflammation in lung epithelial cells. Moreover, they effectively suppressed the MyoFB differentiation induced by TGF-β, exhibiting antifibrotic properties for airway remodeling. LipoFB-EVs also promoted the restoration of AT2 stem cells, which is crucial for lung regeneration. Notably, LipoFB-EVs were found to exclusively enhance colony formation in single AT2 cells derived from patients with COPD, emphasizing their particular efficacy in this context. Although the underlying mechanisms may involve multiple components within LipoFB-EVs, our hypothesis centers on the therapeutic effects being mediated by the heterodimerization of LAT1 and 4F2hc, which facilitates leucine uptake. For instance, enhanced leucine uptake into cells has been associated with increased cellular proliferative capacity through the upregulation of amino acid metabolism, contributing to the restoration of AT2 stem cells in COPD. Furthermore, increased leucine uptake leads to augmented ATP production via acetyl-CoA, thereby alleviating oxidative stress^50^. The mitigation of oxidative stress by leucine uptake has been associated with antifibrotic^51^, anti-inflammatory^52^, and anti-aging effects. Furthermore, heightened intracellular leucine levels exhibit antifibrotic effects by impeding the phosphorylation of SMAD2^53^ and inducing the suppression of the senescence-associated secretory phenotype (SASP), consequently leading to anti-aging effects^54^. Although further investigations and experimental validations are warranted, these findings suggest that LAT1 transferred by LipoFB-EVs may serve as the mechanistic basis for the therapeutic efficacy of this modality in COPD. The intricate interplay between leucine uptake, cellular metabolism, and signaling pathways underscores the potential of LipoFB-EVs to address the multifaceted aspects of COPD pathophysiology.

In experimental models mimicking COPD exacerbation and long-term CS exposure, LipoFB-EVs showed promise in alleviating airway remodeling, inflammation, senescence, and alveolar emphysema. In a short-term exacerbation model, LipoFB-EVs effectively counteracted airway inflammation and remodeling induced by combined cigarette smoke and poly(I:C) treatment. This regimen primarily focuses on exacerbated small airway pathogenesis, which closely mirrors the characteristics observed in human COPD-related airway obstruction. These results are consistent with our *in vitro* findings, highlighting the ability of LipoFB-EVs to mitigate airway remodeling, inflammation, and obstructive airway physiology in a model simulating COPD exacerbation due to CS exposure. In a long-term exposure model, these EVs mitigated airway remodeling, cellular senescence, and alveolar emphysema. Importantly, LipoFB EVs significantly increased the number of LAT1^+^ lung epithelial cells, emphasizing their role in the pathophysiology of COPD. Although the three-month LipoFB-EV treatment did not yield a significant improvement in airway obstruction, as evidenced by FEV0.1%, it enhanced the proportion of AT2 cells and fostered their proliferation and differentiation. These findings underscore the therapeutic regenerative effects of LipoFB-EVs in alveolar emphysema. In tandem with *in vitro* validation results, these findings suggest a plausible mechanism of action involving EV-mediated LAT1 transfer. However, the utilization of human EVs in mouse models necessitates careful consideration, addressing concerns regarding histocompatibility and the potential interactions between human miRNA and protein cargos with their mouse counterparts. In summary, our findings underscore the promising therapeutic potential of LipoFB-EVs in mitigating the diverse aspects of COPD pathology induced by both short- and long-term CS exposure. These effects align with their ability to modulate LAT1 expression and promote regenerative responses, emphasizing the potential of LipoFB-EVs as an intervention for COPD treatment.

Our ongoing clinical development efforts are focused on positioning LipoFB-EVs as a potential therapeutic intervention for patients with COPD. The translation of findings from animal models to human clinical settings presents numerous challenges, including safety assessments, intricacies in dosage optimization, nuanced selection of routes of administration, and stringent regulatory approvals, which are indispensable for the clinical implementation of EV-based therapeutics. Particularly noteworthy in the development of LipoFB-EVs is the scrutiny required for LAT1, the updated expression of which has been reported across diverse cancer types with prognostic implications^55^. In normal tissues, LAT1 exhibits minimal expression and is confined primarily to structures such as the blood-brain and blood-placental barriers, thereby manifesting cancer specificity. Given the propensity of COPD to concomitantly increase the risk of lung cancer, a meticulous examination of its tumorigenic potential is paramount, especially in the context of animal experimentation.

This extensive study sheds light on the intricate role of LipoFBs and their derived EVs in COPD pathogenesis, offers promising avenues for therapeutic interventions, and emphasizes the importance of metabolic reprogramming, cellular communication via EVs, as well as their regenerative potential in lung diseases like COPD. It is, however, important to acknowledge the limitations of this study. First, the potential cross-species interactions between human EVs and their murine counterparts raise concerns about the functional compatibility of human LAT1 (hLAT1) cargos with their murine equivalents. This issue underscores the need to investigate whether the cargo transported by hLAT1 can effectively and functionally engage with murine LAT1 (mLAT1), highlighting a critical aspect in understanding the interplay between human and murine biological systems. Nevertheless, existing knowledge suggests that mLAT1 and hLAT1 exhibit remarkably similar functions as amino acid transporters^56,57^. Previous reports have indicated a comparable inhibition profile of leucine transport using BCH for both mLAT1 and hLAT1^56^. Furthermore, hLAT1 expression in mouse cells has been shown to result in confirmed transport functionality^57^. Taken together, these findings underscore the functional similarities in amino acid transport and inhibition profiles between mLAT1 and hLAT1. Nonetheless, the potential nuances in amino acid transport warrant careful consideration, particularly in the context of using hLAT1-expressing EVs in murine models. Secondly, our study lacked a mechanistic analysis of lung immune cells in a COPD model. The earliest abnormalities induced by smoking in COPD lungs involved hyperplasia of airway basal cells and stem/progenitor cells for ciliated and secretory cells crucial to pulmonary host defense^58^. In contrast, the key inflammatory cells in COPD include macrophages, neutrophils, dendritic cells, and CD8^+^ T cells^59^. While our study observed a reduction in inflammatory cells in the BAL and an inhibitory effect on cytokines due to LipoFB-EVs in a COPD model exacerbated by CS exposure, it is essential to investigate the direct effects on resident cells of the airways, such as alveolar macrophages and neutrophils. Lipid metabolism plays a crucial role in controlling macrophage function, suggesting that LipoFB-EVs have the potential to regulate signal transduction and gene expression during macrophage activation^60,61^. This additional exploration could shed light on the broader impact of LipoFB-EVs on lung immune cells, particularly alveolar macrophages, and contribute to a more comprehensive understanding of their therapeutic potential in COPD. Third, EVs harbor a diverse range of cargo, encompassing proteins, DNA, and RNA, thus contributing to intercellular communication^62^. In this study, our focus was directed toward LAT1 within LipoFB-EVs to evaluate their therapeutic effect on COPD pathology. While assessing the therapeutic potential of LipoFB-EVs, it is challenging to disregard the involvement of cargo components other than LAT1. A comprehensive analysis of miRNAs within LipoFB-EVs revealed their significant role in regulating signals in recipient cells. Unfortunately, despite the profiling of miRNAs in both LF-EVs and LipoFB-EVs, no conspicuous changes were observed, and miRNAs demonstrating therapeutic effects on COPD pathology were not identified (data not shown). Although this study suggests the involvement of LAT1 and 4F2hc in mediating the effects of LipoFB-EVs, the exact mechanisms underlying the observed therapeutic actions remain incompletely understood. Further elucidation of these mechanisms is crucial for establishing robust therapeutic strategies. Addressing these limitations would strengthen the robustness and reliability of the findings and ensure a more comprehensive understanding of the therapeutic potential of LipoFB-EVs in COPD treatment.

In summary, this study paves the way for a more profound understanding of the intricate cellular interactions underlying COPD pathology, potentially opening new avenues for innovative therapeutic interventions. This study delves into a single-cell transcriptome analysis of LipoFBs in COPD-affected lungs, highlighting metabolic dysregulation, notably in the glucose, lipid, and amino acid pathways. Furthermore, EVs derived from LipoFBs demonstrated the potential to suppress epithelial senescence and inflammation, concurrently fostering the restoration of AT2 stem cells. These findings highlight the pivotal role of AT2 cells in maintaining lung health and influencing COPD progression, thereby suggesting that LipoFB(R)-EVs may contribute to their restoration. This groundbreaking approach, harnessing EVs containing secretory factors from LipoFBs, targets the restoration of AT2 stemness and presents a novel therapeutic avenue addressing various facets of COPD pathology, offering promise for preserving the equilibrium of lung airways.

## Acknowledgement

This work was partially supported by Japan Agency for Medical Research and Development, AMED (JP21ym0126007, JP21cm0106402, and JP22ym0126096), Japan Society for the Promotion of Science KAKENHI (JP19K17649, JP21H02930) to Yu Fujita, KAKENHI (JP21H03365) to Shushi Nagamori, and KAKENHI (JP22H03082) to Jun Araya.

## Author contributions

Conceptualization: Yu Fujita; Investigation: Shota Fujimoto, Yuta Hirano, Naoaki Watanabe; Data curation: Shota Fujimoto, Yu Fujita, Methodology: Shota Fujimoto, Yu Fujita,; Software: Shota Fujimoto, Yuta Hirano, Naoaki Watanabe; Formal analysis: Shota Fujimoto, Yuta Hirano, Naoaki Watanabe; Resources: All authors; Writing - Original Draft: Shota Fujimoto, Yu Fujita; Writing - Review & Editing: All authors; Visualization: Shota Fujimoto; Funding acquisition: Yu Fujita, Shushi Nagamori, Jun Araya; Supervision: Yu Fujita; Project administration: Yu Fujita.

