## Supplementary Information for "Lipogenic Lung Fibroblast-derived Extracellular Vesicles Mitigate Cigarette Smoke-Induced Chronic Obstructive Pulmonary Disease Pathologies through LAT1-mediated Alveolar Type II Cell Restoration"

Supplementary Figure 1. Variations in the induction of lipogenic differentiation across LF donor backgrounds.

Supplementary Figure 2. The characterization of LF-EVs and LipoFB-EVs isolated by ultracentrifugation.

Supplementary Figure 3. LipoFB-EVs inhibit the DNA damage response and exhibit superior antifibrotic activity compared to competitor developments.

Supplementary Figure 4. The effects of LipoFB-EVs on HBECs in 2D or 3D cultures.

Supplementary Figure 5. Comparative protein expression between LF-EVs and LipoFB(R)-EVs and the assessment of identified LAT1 Functions.

Supplementary Figure 6. Effects of LipoFB-EVs in a cigarette smoke exposure coupled with poly(I:C) treatment mouse model of COPD.

Supplementary Figure 7. Analysis of mouse lung function and body weight changes in a long-term cigarette smoke-induced mouse model.

Supplementary Table 1：A list of public datasets.

Supplementary Table 2：List of proteins identified by LC-MS/MS in LFs and LipoFBs(R).

Supplementary Table 3：List of proteins identified by LC-MS/MS in LF-EVs and LipoFB(R)-EVs

Supplementary Table 4：List of metabolic pathways included in Reactome Pathway Database

Supplementary Table 5 : Primer information

### Supplementary Figure1

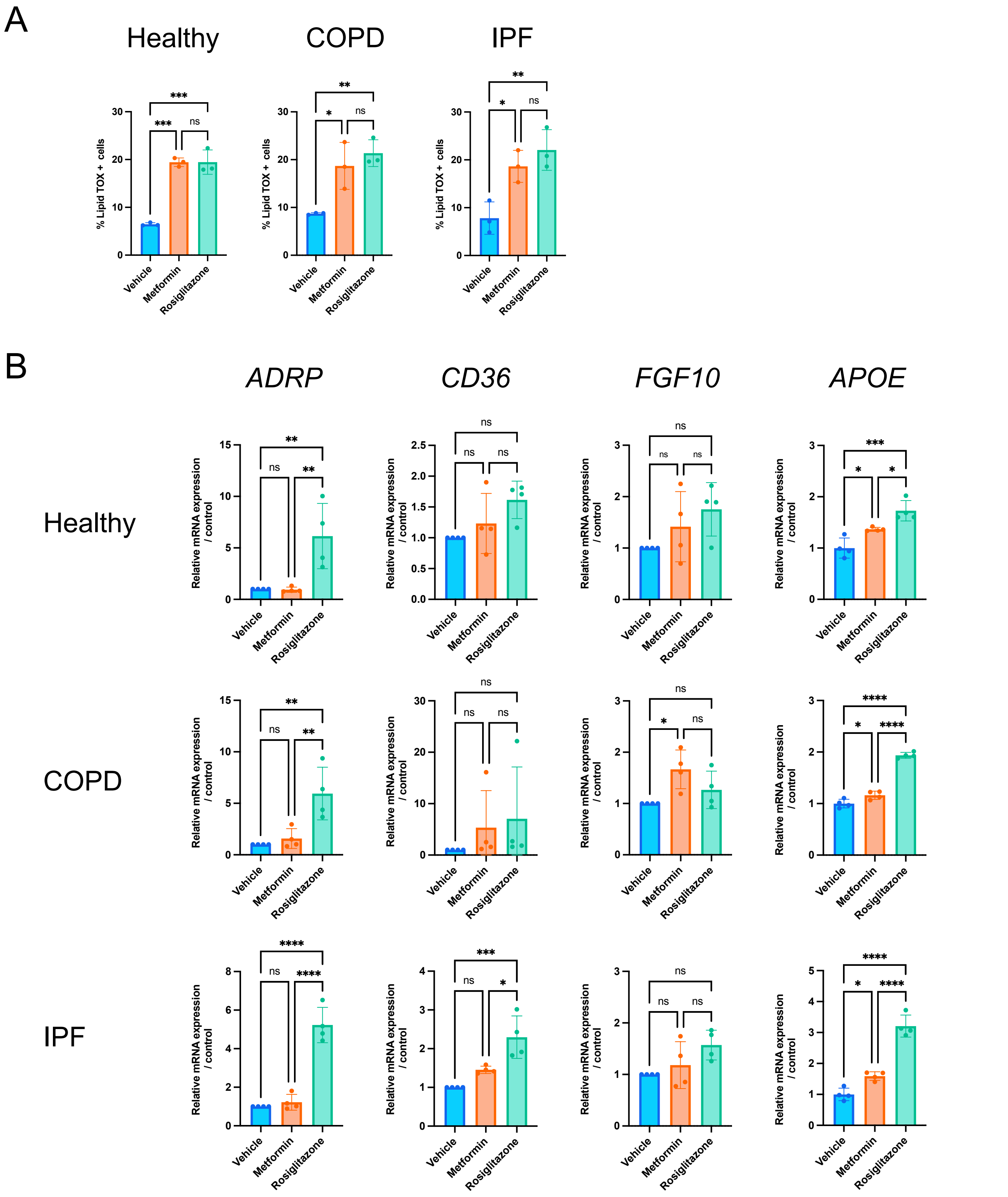


**Supplementary Figure 1. Variations in the induction of lipogenic differentiation across LF donor backgrounds.**

(**A**) Flow cytometry-based quantification of LipidTOX⁺ cell abundance in response to Metformin or Rosiglitazone treatment in healthy subjects, COPD patients, and IPF patients, respectively. ****P* <0.001, ***P* <0.01, **P* <0.05. ns; not significant. (**B**) qPCR analysis for the lipogenic marker genes *ADR*P, *CD36*, *FGF10*, and *APOE* in LFs treated with Metformin, Rosiglitazone, or vehicle in healthy subjects, COPD patients, and IPF patients, respectively. *****P* <0.0001, ****P* <0.001, ***P* <0.01, **P* <0.05. ns; not significant.

Supplementary Figure2

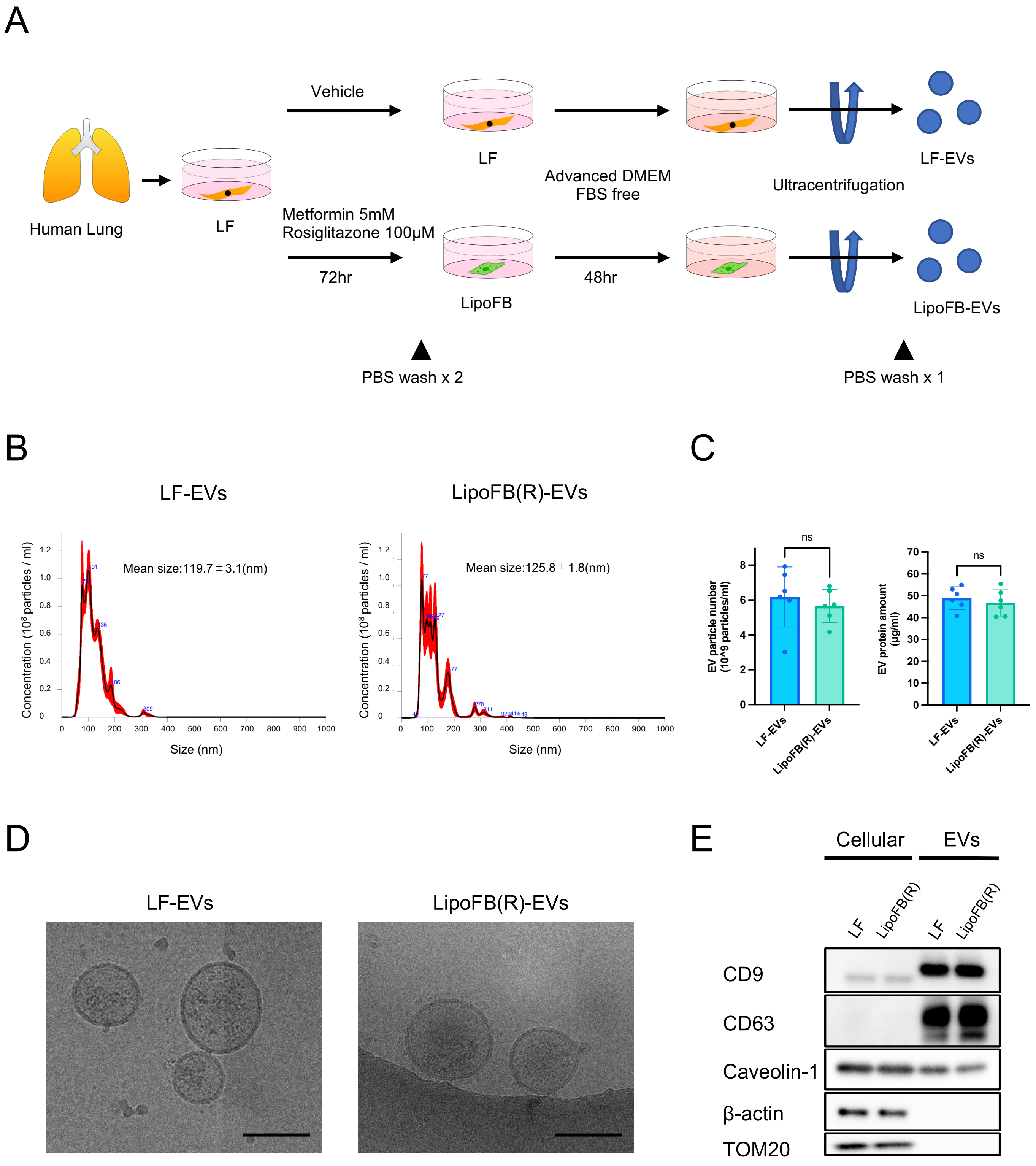


**Supplementary Figure 2. The characterization of LF-EVs and LipoFB-EVs isolated by ultracentrifugation.**

(**A**) Schematic representation of EV purification from the conditioned medium of LFs or LipoFBs by ultracentrifugation. (**B**) Nanoparticle tracking analysis showing particle number and size of LF-EVs and LipoFB(R)-EVs. (**C**) Comparison of EV particle numbers and protein quantity in LF-EVs and LipoFB(R)-EVs. (**D**) Cryo-transmission electron microscopic images in LF-EVs and LipoFB(R)-EVs. (**E**) Representative immunoblot of conventional EV markers for whole cell lysates and EVs from LFs and LipoFB(R).

### Supplementary Figure3

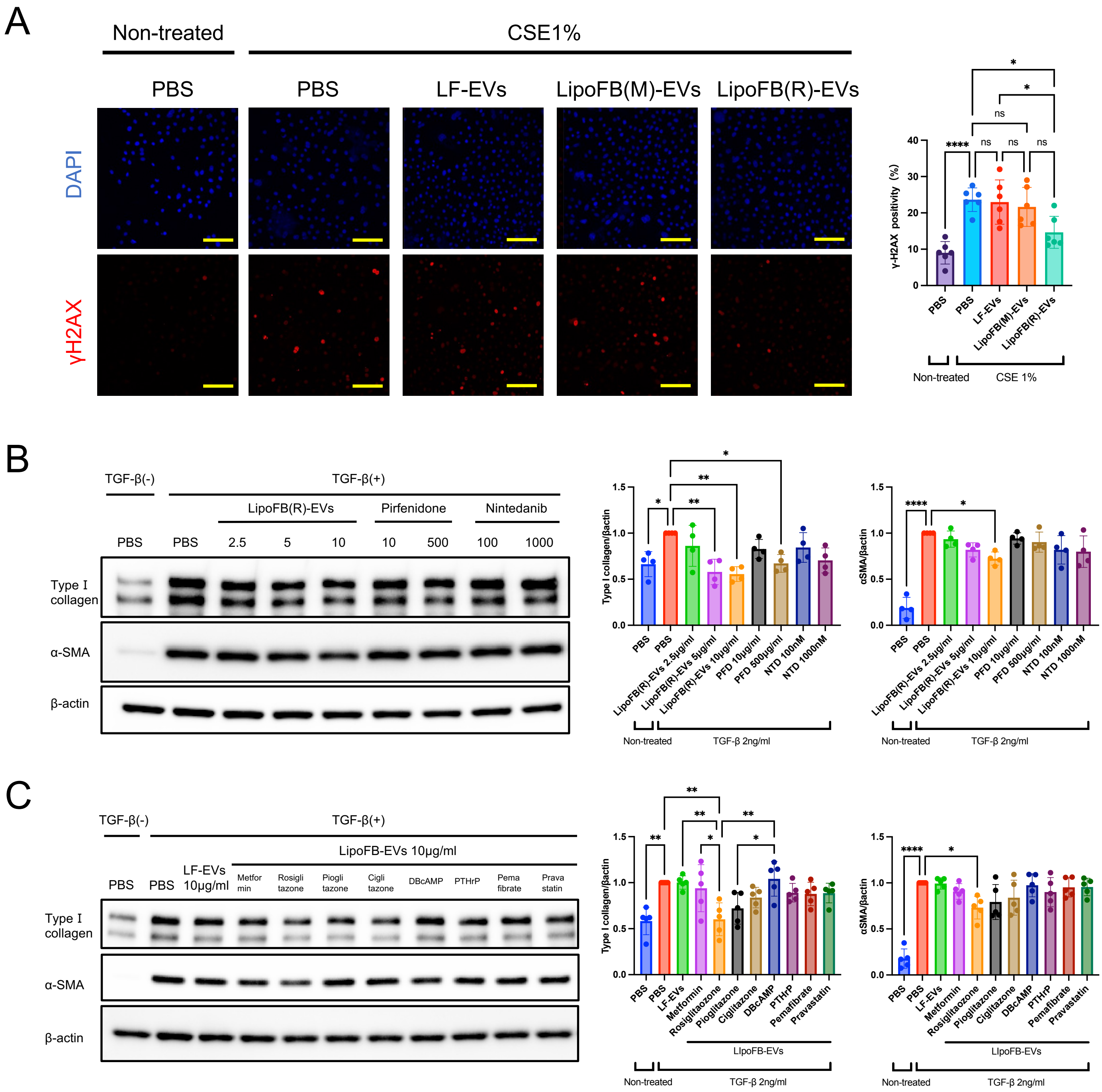


**Supplementary Figure 3. LipoFB-EVs inhibit the DNA damage response and exhibit superior antifibrotic activity compared to competitor developments.**

(**A**) Photographs and quantitative analysis of immunofluorescent staining of phospho-histone H2A.X (Ser139). HBECs were treated for 48 h with PBS, LF-EVs, LipoFB(M)-EVs, or LipoFB(R)-EVs (10 µg/ml) in the presence or absence of CSE (1.0%). ****$P$ <0.0001, *$P$ <0.05. ns; not significant. Scale bars = 100 µm. (**B**) Representative immunoblot and quantitative analysis showing the amount of type I collagen, α-SMA, and β-actin in LFs treated for 24 h with LipoFB(R)-EVs (2.5, 5, or 10 µg/ml), Pirfenidone (10 or 500 µg/ml) or Nintedanib (100 or 1000 nM) in the presence or absence of TGF-β (2 ng/ml for 24h). ****$P$ <0.0001, **$P$ <0.01, *$P$ <0.05. $P$-values are not stated if not significant. (**C**) Representative immunoblot and quantitative analysis showing the amount of type I collagen, α-SMA, and β-actin in LFs treated for 24 h with PBS, vehicle, or EVs derived from various metabolic drug-treated LFs in the presence or absence of TGF-β (2 ng/ml for 24h). Before collecting the conditioned medium form LFs, the LFs were treated by Metforimin, Rosiglitazone, Pioglitazone, Ciglitaozne, DBcAMP, PTHrP, Pemafibrate, or Pravastatin at a concentration of 10µg/ml. ****$P$ <0.0001, **$P$ <0.01, *$P$ <0.05. $P$-values are not stated if not significant.

Supplementary Figure4

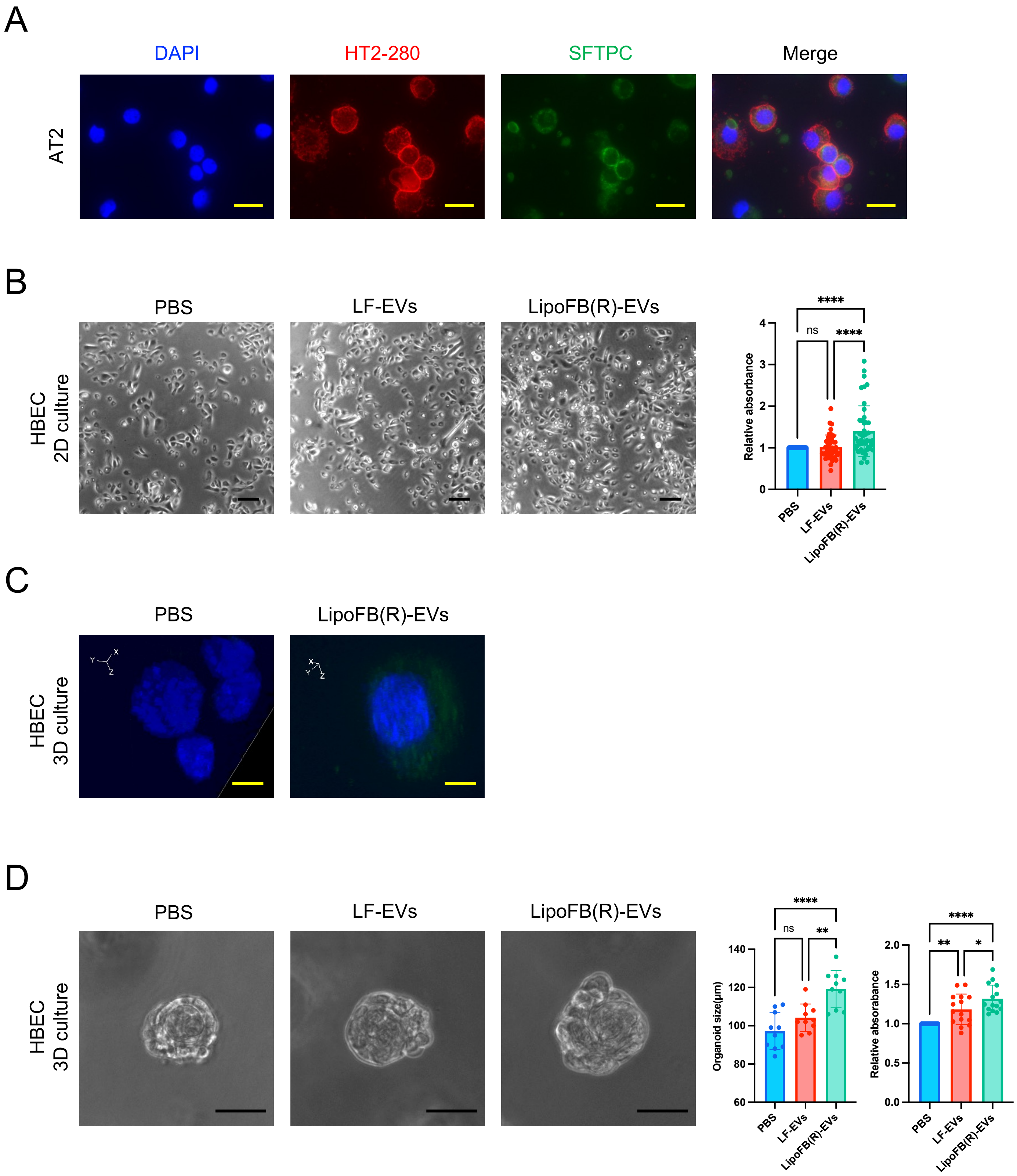


**Supplementary Figure 4. The effects of LipoFB-EVs on HBECs in 2D or 3D cultures.**
(**A**) Validation of human AT2 cells purity using cytospin preparations of HTII-280$^{+}$ sorted cells and staining with HTII-280 (red) and SFTPC (green). Scale bars = 20 μm. (**B**) Representative images and cell counting kit 8 (CCK-8) assay of HBECs in 2D cultures treated with PBS, LF-EVs, or LipoFB-EVs. ****$P$ <0.0001. ns; not significant. Scale bars = 100 μm. (**C**) Confocal microscopy z-stack images of HBECs in 3D cultures incubated with PKH67-labeled LipoFB(R)-EVs (green). Scale bars = 20 μm. (**D**) Representative images, cell counting kit 8 (CCK-8) assay, and organoid size of HBECs in 3D cultures treated with PBS, LF-EVs, or LipoFB(R)-EVs. ****$P$ <0.0001, **$P$ <0.01, *$P$ <0.05. ns; not significant. Scale bars = 50 μm.

### Supplementary Figure5

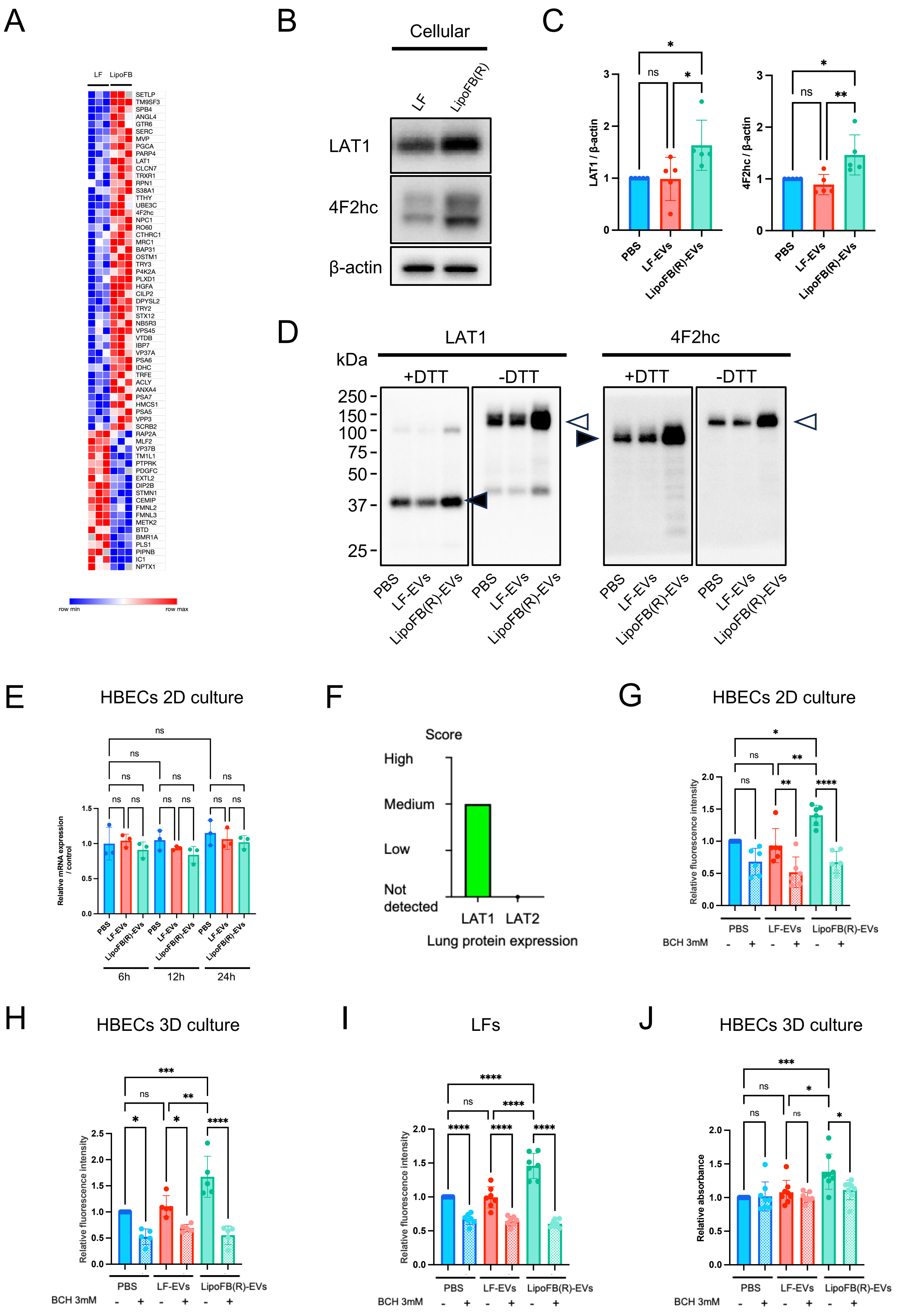

**Supplementary Figure 5. Comparative protein expression between LF-EVs and LipoFB(R)-EVs and the assessment of identified LAT1 Functions.**

(**A**) Heat map of differentially expressed proteins identified by LC-MS/MS between LF-EVs and LipoFB(R)-EVs. Each row in the figure represents a protein, each column is a sample, and the colors represent different expression levels. (**B**) Representative immunoblot of LAT1, 4F2hc, and β-actin in LFs and LipoFB(R) cell lysate. (**C**) Quantitative analysis of LAT1 / β-actin and 4F2hc / β-actin in AT2 cell lysate treated with PBS, LF-EVs, or LipoFB(R)-EVs. (**D**) Expression of LAT1 (left panel) and 4F2hc (right panel) in LFs and LipoFB(R) cell lysate. The cell lysates were treated with (+DTT) or without (−DTT) 100 mM DTT for reducing or non-reducing condition, respectively. The filled arrowhead in the left panel indicates LAT1 monomer. The filled arrowhead in the right panel indicates 4F2hc monomer. The brank arrowheads in the left and right panels indicate heterodimeric complexes of LAT1 and 4F2hc. (**E**) qPCR analysis for relative mRNA expression level of LAT1 in HBECs in 2D cultures treated with PBS, LF-EVs, or LipoFB(R)-EVs at 6, 12, and 24 hr after treatment. ns; not significant. (**F**) The graph depicts the expression patterns of LAT1 and LAT2 proteins in human lung tissues. Data were obtained from the Human Protein Atlas (https://www.proteinatlas.org). (**G**) Relative fluorescence intensity of amino acid uptake assays and inhibition experiments with BCH (3 mM) in HBECs in 2D cultures treated with PBS, LF-EVs, or LipoFB(R)-EVs under sodium-free conditions. ****$P$ <0.0001, **$P$ <0.01, *$P$ <0.05. ns; not significant. (**H**) Relative fluorescence intensity of amino acid uptake assays and inhibition experiments with BCH (3mM) in HBECs in 3D cultures treated with PBS, LF-EVs, or LipoFB(R)-EVs under sodium-free conditions. ****$P$ <0.0001, ***$P$ <0.001, **$P$ <0.01, *$P$ <0.05. ns; not significant. (**I**) Relative fluorescence intensity of amino acid uptake assays and inhibition experiments with BCH (3 mM) in LFs treated with PBS, LF-EVs, or LipoFB(R)-EVs under sodium-free conditions. ****$P$ <0.0001. ns; not significant. (**J**) Inhibition assay using BCH (3 mM) for cell viability of HBECs in 3D cultures. Cell counting kit 8 (CCK-8) assays of HBECs in 3D cultures treated with PBS, LF-EVs, or LipoFB(R)-EVs. ***$P$ <0.001, *$P$ <0.05. ns; not significant.

### Supplementary Figure6

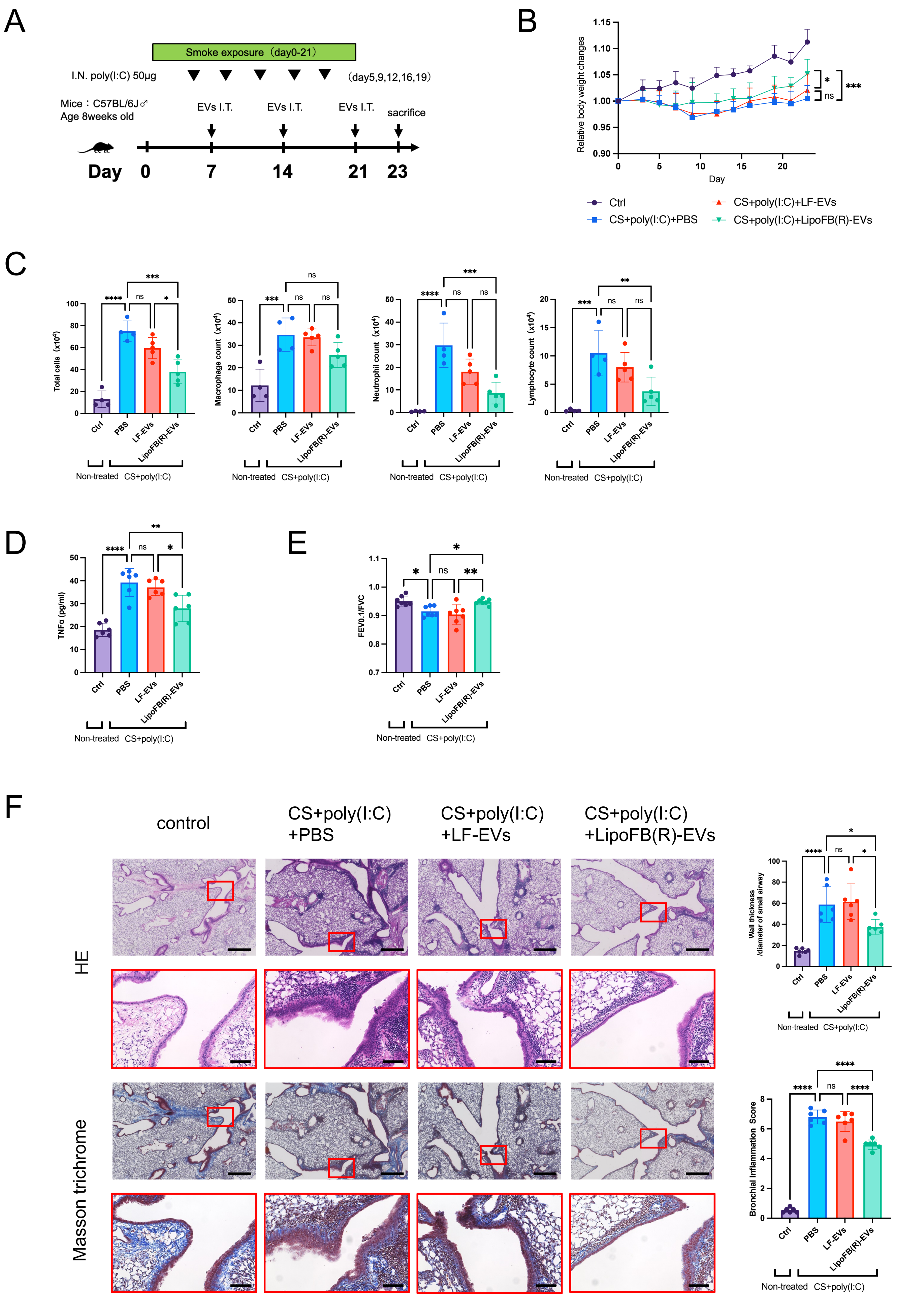

**Supplementary Figure 6. Effects of LipoFB-EVs in a cigarette smoke exposure coupled with poly(I:C) treatment mouse model of COPD.**

(**A**) Schematic protocol for EV treatment in the mouse model of COPD that combined cigarette smoke (CS) with poly(I:C). I.N.: intranasal, I.T.: intratracheal. (**B**) Body weight (BW) changes after CS with poly(I:C) treatment. BW at day 0 before initiating treatment was given a relative value of 1.0. Graphed values represent the mean ± SEM. Control: n=12, CS+poly(I:C)+PBS: n=12, CS+poly(I:C)+LF-EVs: n=12, CS+poly(I:C)+LipoFB(R)-EVs: n=13. (**C**) Cell counts of total cells, macrophages, neutrophils, and lymphocytes in bronchoalveolar lavage fluid (BALF) from mice. Control: n=4, CS+poly(I:C)+PBS: n=4, CS+poly(I:C)+LF-EVs: n=5, CS+poly(I:C)+ LipoFB(R)-EVs: n=5. ****$P$ <0.0001, ***$P$ <0.001, **$P$ <0.01, *$P$ <0.05. ns; not significant. (**D**) TNF-α expression was detected in lung tissue by ELISA. Control: n=6, CS+poly(I:C)+PBS: n=6, CS+poly(I:C)+LF-EVs: n=6, CS+poly(I:C)+ LipoFB(R)-EVs: n=6. ****$P$ <0.0001, **$P$ <0.01, *$P$ <0.05. ns; not significant. (**E**) FEV0.1/FVC ratio was measured in mice. Parameters are shown for each individual mouse, along with group averages (±SEM). Control: n=7, CS+poly(I:C)+PBS: n=7, CS+poly(I:C)+LF-EVs: n=7, CS+poly(I:C)+ LipoFB(R)-EVs: n=7. **$P$ <0.01, *$P$ <0.05. ns; not significant. (**F**) Low and high magnification images of histology of representative lung sections from each group by haematoxylin and eosin (HE) staining and Masson's trichrome staining. Quantification of small airway wall thickness and inflammation score. Scale bar for weakly magnified images is 500 μm; scale bar for strongly magnified images is 100 μm. Control: n=6, CS+poly(I:C)+PBS: n=6, CS+poly(I:C)+LF-EVs: n=6, CS+poly(I:C)+ LipoFB(R)-EVs: n=6. ****$P$ <0.0001, *$P$ <0.05. ns; not significant.

### Supplementary Figure7

A

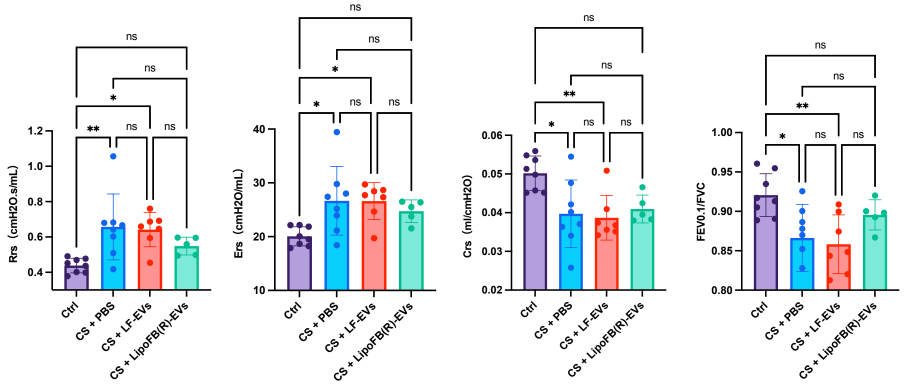


B

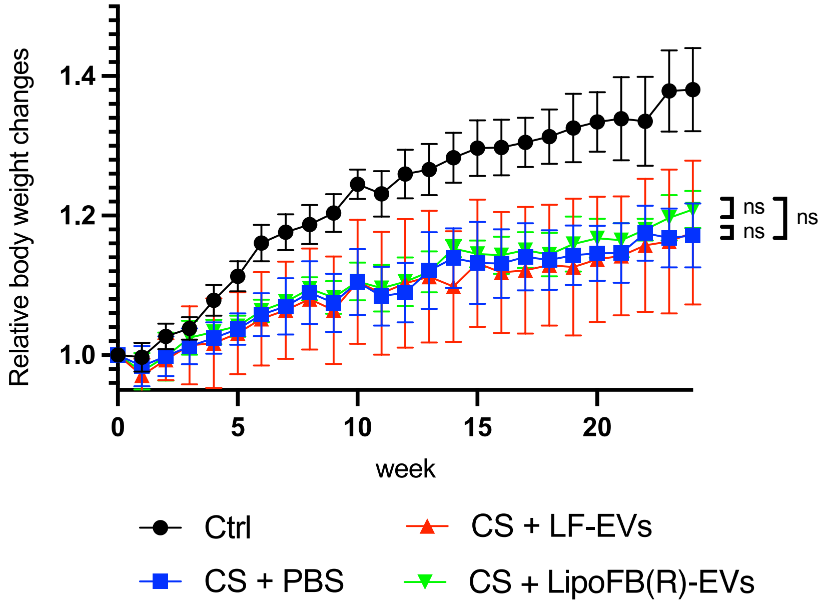


**Supplementary Figure 7. Analysis of mouse lung function and body weight changes in a long-term cigarette smoke-induced mouse model.**

(**A**) flexiVent analysis of airway resistance (Rn), elastance (Ers), dynamic compliance (Crs), and percent forced expiratory volume in 0.1 second (FEV0.1%: FEV0.1/FVC). ****P* <0.01, **P* <0.05. ns; not significant. n=8 in the control group, n=8 in the CS+PBS group, n=7 in the CS+LF-EVs group, and n=5 in the CS+LipoFB(R)-EVs group. (**B**) Body weight (BW) changes after CS treatment. BW at day 0 before initiating treatment was given a relative value of 1.0. Graphed values represent the mean ± SEM. n=8 in the control group, n=8 in the CS+PBS group, n=7 in the CS+LF-EVs group, and n=5 in the CS+LipoFB(R)-EVs group. ns; not significant.
